# Stress induces the transcription of toxin-antitoxin systems but does not activate toxin

**DOI:** 10.1101/2020.03.02.972737

**Authors:** Michele LeRoux, Peter H. Culviner, Yue J. Liu, Megan L. Littlehale, Michael T. Laub

**Affiliations:** Department of Biology, Massachusetts Institute of Technology, Cambridge, MA 02139, USA; Howard Hughes Medical Institute, Massachusetts Institute of Technology, Cambridge, MA 02139, USA

## Abstract

Toxin-antitoxin (TA) systems are ubiquitous genetic elements in bacterial genomes, but their functions are controversial. Although they are frequently postulated to regulate cell growth following stress, few null phenotypes for TA systems have been reported. Here, we show that TA transcript levels can increase substantially in response to stress, but toxin is not liberated. We find that the growth of an *Escherichia coli* strain lacking 10 TA systems encoding endoribonuclease toxins is not affected following exposure to six stresses that each trigger TA transcription. Additionally, using RNA-sequencing, we find no evidence of mRNA cleavage following stress. Stress-induced transcription arises from antitoxin degradation and relief of transcriptional autoregulation. Importantly, although free antitoxin is readily degraded *in vivo*, antitoxin bound to toxin is protected from proteolysis, preventing release of active toxin. Thus, transcription is not a reliable marker of TA activity, and TA systems likely do not strongly promote survival following stress.

## Introduction

Toxin-antitoxin (TA) systems are composed of a growth-inhibiting toxin and a cognate, neutralizing antitoxin. These systems are found in the majority of bacterial chromosomes and also on plasmids, phage, and other mobile elements, with many species encoding dozens of different systems. Despite their prevalence, the functions of most TA systems remain unknown. They have been suggested to promote stress tolerance, genome stability, persister cell formation, and programmed cell death, but the evidence supporting these functions is often limited, speculative, or controversial (Christensen et al., 2003; Gerdes and Maisonneuve, 2012; Harms et al., 2016; Hazan et al., 2004; Kolodkin-Gal and Engelberg-Kulka, 2006; Magnuson, 2007; Nigam et al., 2019; Ronneau and Helaine, 2019; Van Melderen and Wood, 2017; Wang et al., 2011; Yamaguchi et al., 2011).

TA systems are classified by the nature of the antitoxin, with the most well-characterized versions, termed type II systems, consisting of a protein that binds the cognate toxin to prevent its activity. These type II toxin-antitoxin pairs are encoded in the same operon and co-expressed (Fig. 1A). The targets of toxins from type II systems vary, but most inhibit a central cellular process such as translation or DNA replication (Yamaguchi et al., 2011). The direct targets have been identified by ectopically expressing individual toxins, which are often bacteriostatic and thus reversible when antitoxin levels are restored. Most antitoxins are small proteins that harbor unstructured or disordered regions and are thought to be substrates of proteases such as Lon and ClpP (Gerdes and Maisonneuve, 2012; Yamaguchi et al., 2011). Although toxin-antitoxin pairs are co-transcribed, antitoxins are usually translated at a higher rate, presumably to outpace antitoxin protein turnover and maintain sufficient levels for neutralizing toxin (Li et al., 2014). In addition to binding their cognate toxins, most antitoxins transcriptionally autoregulate by directly binding the TA operon promoter to repress transcription. Many TA systems exhibit so-called conditional cooperativity, in which low concentrations of toxin promote the binding of antitoxin to its promoter whereas higher concentrations of toxin disrupt binding (Overgaard et al., 2008).

**Figure 1.**
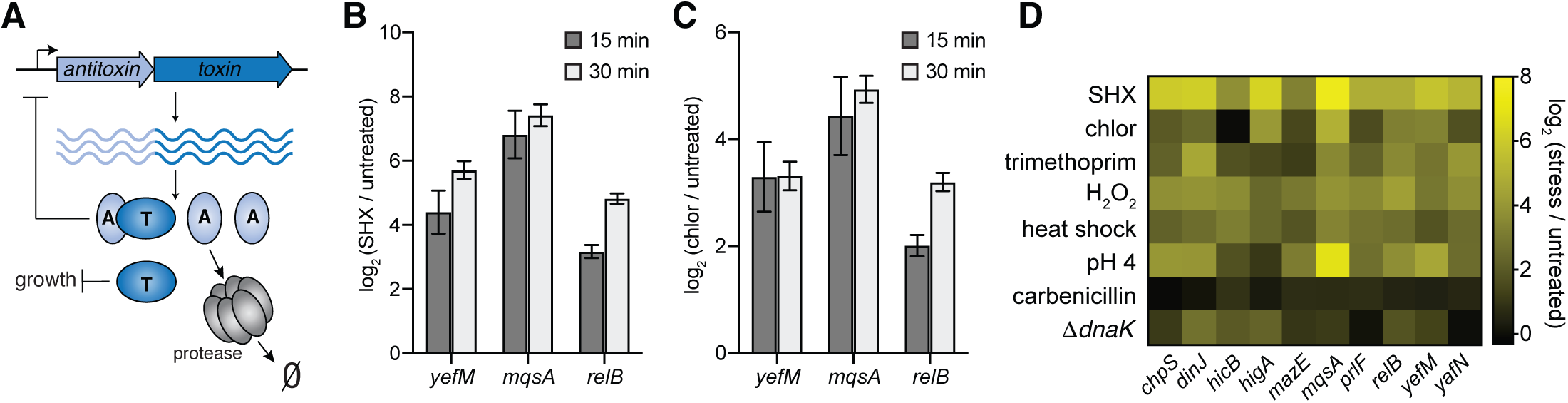
TA systems are transcriptionally induced by diverse stresses. (A) Schematic summarizing key properties of toxin-antitoxin systems. Toxin and antitoxin are co-transcribed from a single operon, but the antitoxin is translated at a higher rate. Antitoxins bind to and repress their own promoters. Antitoxins are also substrates of cellular proteases such as Lon. (B-C) Changes in antitoxin transcript level in response to serine hydroxamate (SHX) (B) or chloramphenicol (chlor) (C) as measured by qRT-PCR at the indicated time points. Antitoxin levels were normalized to the housekeeping gene *gyrA* and each stress normalized to measurements made for untreated cells. (D) Difference in antitoxin transcript levels between cells exposed to stress for 30 minutes compared to untreated cells as measured by qRT-PCR. Data are the average of 3 biological replicates with error bars indicating the S.D. Also see Figure S1.

The first type II TA system identified, CcdAB, is plasmid-encoded and was found to promote plasmid inheritance through post-segregational killing (Jaffé et al., 1985; Salmon et al., 1994). The antitoxin CcdA is both highly expressed and intrinsically unstable (Van Melderen et al., 1994). Thus, if the plasmid harboring *ccdAB* is not segregated into a daughter cell, CcdA is degraded and cannot be replenished, leading to liberation of the CcdB toxin which can kill cells, thereby ensuring that plasmid-free cells do not survive.

Type II TA systems were subsequently found encoded on bacterial chromosomes. These systems do not stabilize chromosomes as with CcdAB, raising the question of what function, if any, they provide cells. Chromosomal TA systems are typically assumed to have similar properties to CcdAB. However, antitoxin degradation rates have, to our knowledge, never been examined in their native contexts using pulse-chase experiments. Instead, antitoxin degradation is most commonly examined by high ectopic expression coupled with translation-shutoff experiments or by *in vitro* degradation assays (Cherny, 2005; Christensen et al., 2001; Prysak et al., 2009; Wang et al., 2011). Many studies have invoked TA systems as stress response elements, primarily because they are often transcriptionally upregulated in stressful conditions, including nutritional limitations, heat shock, and oxidative stress (Christensen-Dalsgaard et al., 2010; Muthuramalingam et al., 2016; Ronneau and Helaine, 2019). However, increased transcription of a TA system does not necessarily imply that toxin is liberated and active. Notably, there are very few reported cases of chromosomal TA systems that have null phenotypes in the stress conditions that lead to their increased transcription.

An early, influential study on the RelBE TA system found that transcription increased following nutritional stress and that global cellular translation rates were ∼two-fold higher in a Δ*relBE* strain compared to the wild type in those same conditions, but the reproducibility of this small effect is unclear and no growth phenotype was reported for the Δ*relBE* strain (Christensen and Gerdes, 2004). A subsequent study examined a strain lacking five TA systems in *E. coli*, finding that it did not grow better following several different stresses (Tsilibaris et al., 2007). A study of the YefM-YoeB system found that its toxin, YoeB, cleaves an ectopically expressed, artificial mRNA featuring multiple ribosome pause sites following heat shock, but did not see cleavage of select native transcripts or a YoeB-dependent growth phenotype (Janssen et al., 2015).

A potential breakthrough came when the Gerdes lab reported that the sequential deletion of 10 chromosomal TA systems in *Escherichia coli* (Δ10TA) significantly reduced the frequency of antibiotic-tolerant persister cells (Maisonneuve et al., 2011). Their subsequent work argued that guanosine penta- and tetra-phosphate, (p)ppGpp, the master regulator of the bacterial stringent response, induces TA systems by driving the accumulation of polyphosphate to stimulate the Lon protease to degrade antitoxins at an accelerated rate, thereby freeing toxins to promote persister cell formation (Maisonneuve et al., 2013). However, much of this work was retracted when it came to light that the Δ10TA strain had acquired multiple ϕ80 prophage insertions that were responsible for the persister phenotypes observed (Goormaghtigh et al., 2018; Harms et al., 2017).

Since the discovery of chromosomal TA systems 30+ years ago, numerous studies have invoked a role in responding to stress, but there is little evidence that these systems alter cellular physiology during or after stress. To address this paradox, we set out to understand (i) how TA transcription increases following abiotic stress and (ii) whether these increases in transcription are associated with toxin activity. We determined that six diverse stresses induce TA transcription, often quite substantially. However, we find that a Δ10TA strain (lacking ϕ80 prophages) does not grow any faster than a wild-type strain following these stresses as would be the case if stress activated any of these growth-inhibitory toxins. Additionally, using RNA-seq, we find no evidence of endoribonuclease toxin activity following stress. Through detailed study of two *E. coli* K12 TA systems, YefM-YoeB and MqsA-MqsR, we find that increases in TA transcription following stress arise from a relief of autoregulation. Stress triggers either a decrease in synthesis of toxin and antitoxin coupled to ongoing degradation of antitoxins, or it accelerates antitoxin degradation, though not in a ppGpp-dependent manner, as previously suggested (Christensen et al., 2001; Maisonneuve and Gerdes, 2014). In either case, antitoxin turnover relieves autoregulation, leading to transcriptional induction. Importantly, we demonstrate that free antitoxin is preferentially degraded relative to antitoxin in complex with toxin. Consequently, toxin remains inhibited and is not liberated. In sum, our results strongly suggest that although TA systems are transcriptionally induced by stress, they are likely not critical effectors of bacterial stress responses, as is frequently asserted.

## Results

### Type II TA systems are transcriptionally induced by diverse stress conditions

Numerous studies have reported increased transcription of type II TA systems in response to diverse stress conditions. To determine the kinetics of these transcriptional responses, we subjected *E. coli* to two well-studied stresses, the stringent response to amino acid starvation, which can be rapidly induced via addition of serine hydroxamate (SHX), and translation inhibition, induced by treatment with chloramphenicol. Consistent with previous reports (Christensen-Dalsgaard et al., 2010; Muthuramalingam et al., 2016; Ronneau and Helaine, 2019; Shan et al., 2017), we found an increase of at least 6-fold in the mRNA levels of three type II TA systems in *E. coli* MG1655, *mqsRA*, *relBE*, and *yefM-yoeB,* at both 15 and 30 minutes after treatment using quantitative reverse transcription PCR (qRT-PCR) (Fig. 1B-C).

Based on these data, we selected the 30-minute time point for subsequent qRT-PCR experiments, and examined the expression of 10 type II systems of *E. coli* MG1655 in response to a larger panel of stresses including amino-acid starvation (SHX), translation inhibition (chloramphenicol), DNA synthesis inhibition (trimethoprim), oxidative stress (hydrogen peroxide), cell-wall synthesis inhibition (carbenicillin), acid shock (pH 4), heat shock (shift from 30 °C to 45 °C), and proteotoxic stress (Δ*dnaK*) (Fig. 1D). We saw significant increases in antitoxin transcription, in some cases exceeding 50-fold. Although the magnitude of the responses varied across TA systems and not all systems responded to all stresses, each stress except carbenicillin elicited a transcriptional response in the majority of *E. coli* TA systems (Fig. 1D, Fig. S1).

### Type II TA systems do not affect growth following diverse stress conditions

An increase in TA transcription during stress has frequently been interpreted to mean that TA systems are ‘active’ under these conditions. We reasoned that if toxin were released following a stress, we should see a TA-dependent decrease in growth during or immediately after the stress. To test this hypothesis, we used a recently constructed Δ10TA strain in which each of 10 type II TA systems in *E. coli* MG1655 have been deleted to examine growth following a stress (Goeders et al., 2013; Goormaghtigh et al., 2018). This strain was verified as lacking the prophage contaminant found in a different, prior version of this strain. For each stress that induced TA transcription in Figure 1, we grew the wild-type and Δ10TA strains to early exponential phase and then split each culture, treating half with a given stress for one hour while leaving the other half untreated for one hour (Fig. 2A). The cells were then washed to remove the stress agent and diluted into fresh medium. The growth of each culture was then measured for 8-10 hours. Despite the fact that each of these stresses induces TA transcription (Fig. 1D), none of them led to a difference in growth rate between the wild-type and Δ10TA strain for up to 8 hours (Fig. 2B-I). The lag phase of the Δ10TA strain was increased following hydrogen peroxide treatment compared to the wild-type (Fig. 2E); however, this is opposite of what is expected if toxins were liberated, and it may result from another mutation acquired in construction of the Δ10TA strain. To rule out the possibility that small changes in growth were undetectable in these experiments, or that a single stress treatment was insufficient to stimulate detectable toxin activity, we also performed a competition experiment in which differentially labeled wild-type and Δ10TA strains were mixed at a 1:1 ratio and subjected to two cycles of chloramphenicol treatment. The Δ10TA strain displayed no competitive advantage or disadvantage, again indicating that toxins are not liberated by this stress (Fig. 2J).

**Figure 2.**
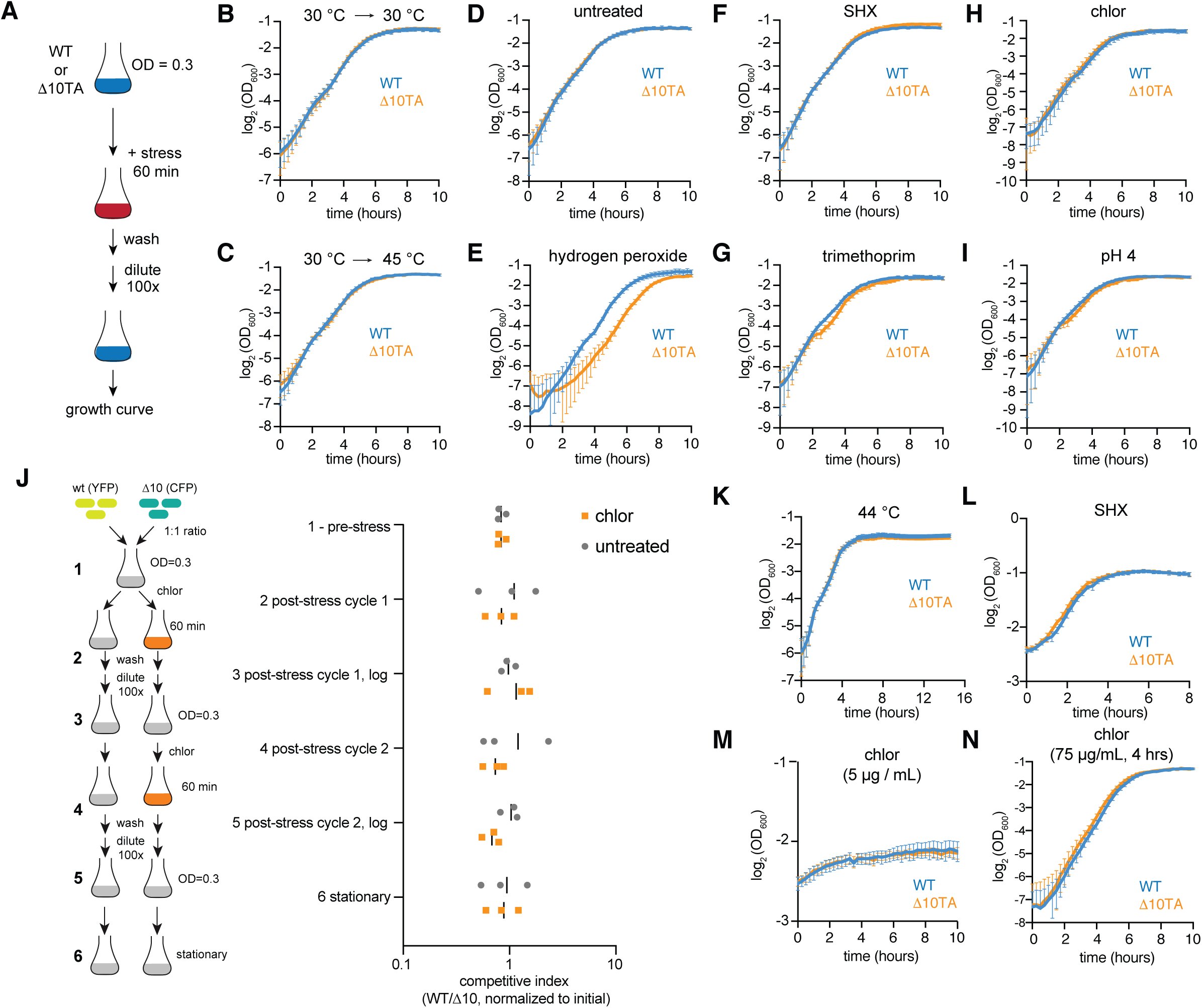
TA systems do not inhibit growth following diverse stress conditions. (A) Experimental design for panels A-J to measure the growth of wild-type and Δ10TA strains following stress exposure. (B-C) Growth curves for WT and Δ10TA strains grown at 30 °C to OD_600_ of ∼0.3, then maintained at 30 °C (B) or shifted to 45 °C (C) for 60 min before dilution and growth rate measurement at 37 °C in flasks. (D-I) Growth curves for WT and Δ10TA strains grown at 37 °C and left untreated (D) or treated with the indicated stress for one hour then washed and diluted into fresh media, as described in (A). For complete description of stress conditions, see Methods. (J) Wild-type and Δ10TA strains labeled with YFP and CFP, respectively, were mixed at a 1:1 ratio. This mixture was propagated and treated with two cycles of chloramphenicol as depicted in the schematic. Samples were taken points indicated by numbers on left, and colony forming units (c.f.u.) of the two strains were determined. The resulting competitive index is graphed on the right where individual symbols indicate independent replicates; black bar represents mean. (K) Growth curve of WT and Δ10TA strains grown at 44 °C. (L-M) Growth curve of WT and Δ10TA strains grown to OD600 ∼0.3, then treated with the indicated stress throughout growth measurements. (N) Growth curves for WT and Δ10TA strains grown at 37 °C and treated with chloramphenicol for four hours, then washed and diluted into fresh media, as described in (A).

To test for differences in growth during the stress itself, we also monitored growth at 44 °C (Fig. 2K), and immediately after adding SHX and chloramphenicol without washing out the stress agent (Fig. 2L-M). In none of these conditions did we observe significant differences in growth between the wild-type and Δ10TA strains. We also examined growth after longer exposure (4 hours) to a growth-inhibitory concentration of chloramphenicol (75 µg/mL) followed by washout of the stressor, and saw a very subtle growth difference (Fig. 2N). Although a minor difference was evident after these extended periods of chloramphenicol stress, TA transcription increased within minutes in the same conditions (Fig. 1B-C). Thus, we conclude that toxins are not strongly or significantly activated by stress and that transcriptional activation is not a reliable proxy for toxin activity.

### RNA sequencing reveals no evidence of toxin activity

For a more direct assay of toxin activity, we turned to RNA sequencing (RNA-seq) as each of the 10 toxins deleted in the Δ10TA strain are endoribonucleases. The endoribonuclease activity of a toxin can be quantitatively measured by comparing RNA-seq fragment-densities between cells expressing a given toxin and cells not expressing the toxin (Culviner and Laub, 2018) (Fig. 3A). A ratio of fragment density +/− a given toxin can be calculated for each nucleotide in each transcript. Regions where the ratio is negative may indicate toxin cleavage, or could reflect changes in transcript levels or stability that arise after toxin expression. To distinguish between toxin activity and other changes to mRNA abundance, we focused on areas with steep “valleys”, which are characteristic of toxin cleavage events (Culviner and Laub, 2018). Because the toxin MazF – one of the toxins deleted in the Δ10 strain – has been extensively characterized and has a well-defined cleavage motif, we focused the first part of our analysis on this toxin. An example profile of fragment density following the induction of MazF for 10 min is shown in Fig. 3A (Culviner and Laub, 2018). This profile exhibits a clear signature of MazF cleavage: a region with a low ratio, ∼30 nucleotides into the *rplS* transcript, followed by a steep increase that coincides with a high-scoring MazF motif (Fig. 3A, bottom). This high positive slope in the ratio profile occurs because MazF typically generates one product that can be rapidly degraded by 3’-5’ exonucleases and one product that is more stable as *E. coli* lacks a 5’-3’ exonuclease.

**Figure 3.**
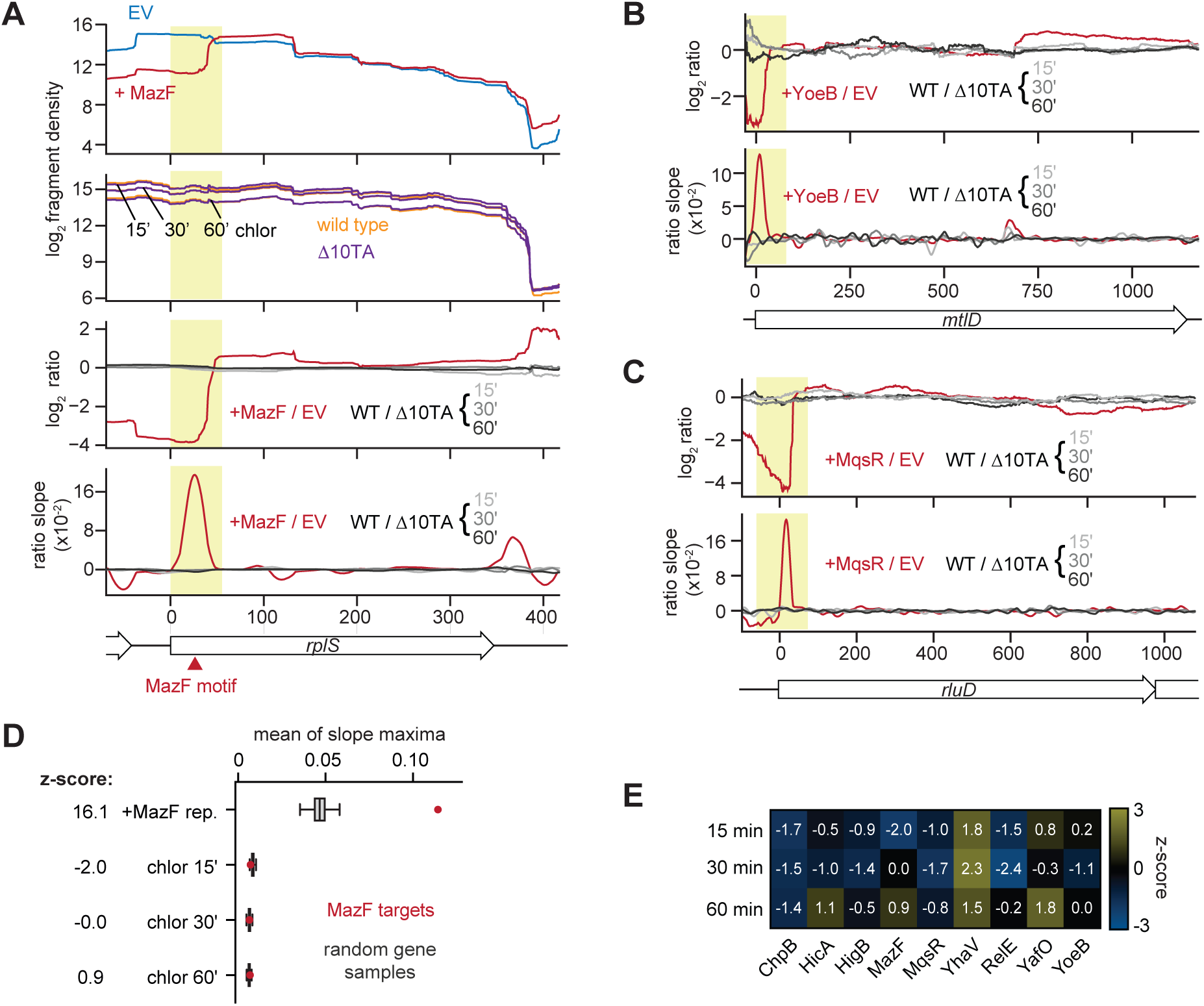
RNA-seq reveals no evidence of endoribonuclease toxin activity following stress. (A) Example analysis for MazF cleavage site (yellow shaded box) found within the *rplS* transcript. Summed read counts for empty vector and MazF overexpression (upper panel) and wild-type or Δ10TA following chloramphenicol treatments at indicated time points (second panel). The ratios of MazF / empty vector and WT / Δ10TA in chloramphenicol are shown (third panel) with the slop of the ratio profiles below (fourth panel). The known MazF cleavage motif is indicated (red arrow). (B-C) Ratio profiles (upper) and resulting ratio slopes (lower) are shown for YoeB cleavage site (yellow box) in the *mtlD* transcript (B) and MqsR cleavage site (yellow box) in the *rluD* transcript (C). (D) Average of the slope maxima in the ratio profiles of coding regions corresponding to ‘MazF targets’ as defined in the text (red dot), compared to a distribution generated from sampling random sets of coding regions of the same size 10,000 times from the same sample (grey box and whiskers). The z-score calculated from this comparison is indicated (left). The same analysis was performed on the WT v. Δ10TA in chloramphenicol data sets, comparing MazF targets to a distribution based on random samples. (E) Z-scores resulting from the analysis described in (D), but for each of the nine TA systems indicated. See also Figures S2 and S3.

We recently generated RNA-seq data for cells expressing each of 9 endoribonuclease toxins in *E. coli* for 5 min, including MazF (see Methods). These toxins are those deleted in the Δ10TA strain; YafQ was excluded as no cleavage or growth defect was detected following its overexpression. To test whether stress induces activity of any the endoribonuclease toxins, we performed RNA-seq on wild-type and Δ10TA cells following 15, 30, and 60 min of exposure to a concentration of chloramphenicol that results in high TA transcription (Fig. 1). We then calculated a ratio of fragment density at each nucleotide in each transcript for wild-type v. Δ10TA cells. These ratios were compared to the ratios generated following the overexpression of each toxin. We examined regions with strong signatures of cleavage following toxin overexpression, but found no evidence of TA-dependent mRNA cleavage following chloramphenicol treatment at any of the time points tested (Fig. 3A-C, Fig. S2A-I). The ratios of fragment density for wild type v. Δ10TA following stress did not exhibit large valleys like those seen after ectopically producing an individual toxin.

We also considered the possibility that stress leads only to very modest cleavage or cleavage in only a subset of cells. If so, we would expect ratios of fragment density for wild type v. Δ10TA to have a similar overall shape as those produced following ectopic toxin expression, but with shallower valleys. For instance, in cells in which MazF expression is titrated to levels that allow continued cell growth (Culviner and Laub, 2018), the +/− toxin profiles retain very similar overall shapes, but with a reduction in magnitude (Fig. S3A). A close inspection of the cleavage profiles following chloramphenicol treatment did not reveal any substantial similarity to the profiles generated after ectopic toxin expression (Fig S3A).

To more systematically assess whether chloramphenicol stress activates the endoribonuclease toxins, we first selected the transcripts containing the largest (top 5%) slopes in the ratio profiles resulting from expression of MazF, thereby defining a set of ‘MazF targets’. We then (i) identified the maximum slope for each of these transcripts in the profiles generated by comparing wild type v. Δ10TA in chloramphenicol and (ii) calculated the mean of these slopes (see red dots in Fig. 3D). We repeated this same process but for 10,000 different, random sets of transcripts to produce a distribution reflecting the expected value of mean slope maxima for a random set of transcripts. We could then calculate a z-score for the mean slope maxima of the MazF targets (Fig. 3D). A large z-score would indicate that MazF targets are cleaved more following chloramphenicol stress than expected by chance. However, the z-scores for MazF targets in the three chloramphenicol data sets were −2.0, 0, and 0.9 for the 15, 30, and 60 minute time-points respectively (Fig. 3D) indicating no significant evidence of cleavage. As a control, we confirmed that the MazF targets do yield a high z-score of 16.1 when compared to a replicate of the MazF expression data rather than chloramphenicol stress (Fig. 3D).

We performed this same overall procedure for the targets of the other 8 endoribonuclease toxins. We did not observe high z-scores for any of the other toxin targets in any of the three chloramphenicol data sets (Fig. 3E). YhaV had the highest z-scores of 1.8, 2.3, and 1.5, but inspection of the profiles for the transcripts defined as YhaV targets did not reveal signatures of cleavage in wild type v. Δ10TA in chloramphenicol, i.e. the shape of these profiles did not resemble those resulting from YhaV induction (Fig. S3B). Taken together, these results support our conclusion that there is no substantial activation of toxins in cells exposed to chloramphenicol for up to 60 minutes.

The analyses above are based on the assumption that large slopes in the profiles generated by expression individual toxins appropriately capture cleavage events. To consider the possibility of cleavage events that do not result in steep slopes (e.g. if the 5’ end created by cleavage is not stable, and leads to a wide valley in the +/− toxin ratio plots), we repeated our analysis using minima within transcripts rather than slopes to define toxin targets; in this analysis, low z-scores would indicate cleavage of toxin targets in the chloramphenicol data sets (Fig. S3D-E). However, the lowest z-score in the chloramphenicol treatment data set was only −1.9, for YafO at 60 min (Fig. S3E).

Taken all together, our results indicate no clear, detectable evidence of cleavage by any endoribonuclease toxin in the wild type relative to the Δ10 strain following 15, 30, or 60 minutes of chloramphenicol stress. These findings support our growth measurements also indicating that toxins are not active following stress, despite their strong transcriptional induction.

### Transcriptional induction results from a relief of autoregulation

We wanted to understand, at a mechanistic level, how TA systems are transcriptionally induced without apparently liberating toxin. We focused on three TA systems: YefM-YoeB, MqsRA, and RelBE. To test whether the stress-induced transcription of these systems stems from a loss of transcriptional autoregulation, we first generated chromosomally-encoded antitoxin alleles with mutations in residues that are required for DNA-binding but that do not affect the ability of the antitoxin to neutralize its cognate toxin: *mqsA(N97A/R101A)*, *relB(R7A)*, and *yefM(R10A)* (Bailey and Hayes, 2009; Brown et al., 2011; Overgaard et al., 2009) hereafter referred to as *mqsA\**, *relB\**, and *yefM\**, respectively. As expected, baseline expression of these transcripts was elevated compared to the wild-type alleles (Fig. 4A). We then tested whether these strains still mounted a transcriptional response to chloramphenicol stress. In each case, the DNA-binding mutant no longer exhibited a large increase in transcription as seen with the wild type (Fig. 4B). These results indicate that the transcriptional responses seen in wild-type cells reflect a relief of autoregulation, likely resulting from an increase in antitoxin degradation.

**Figure 4.**
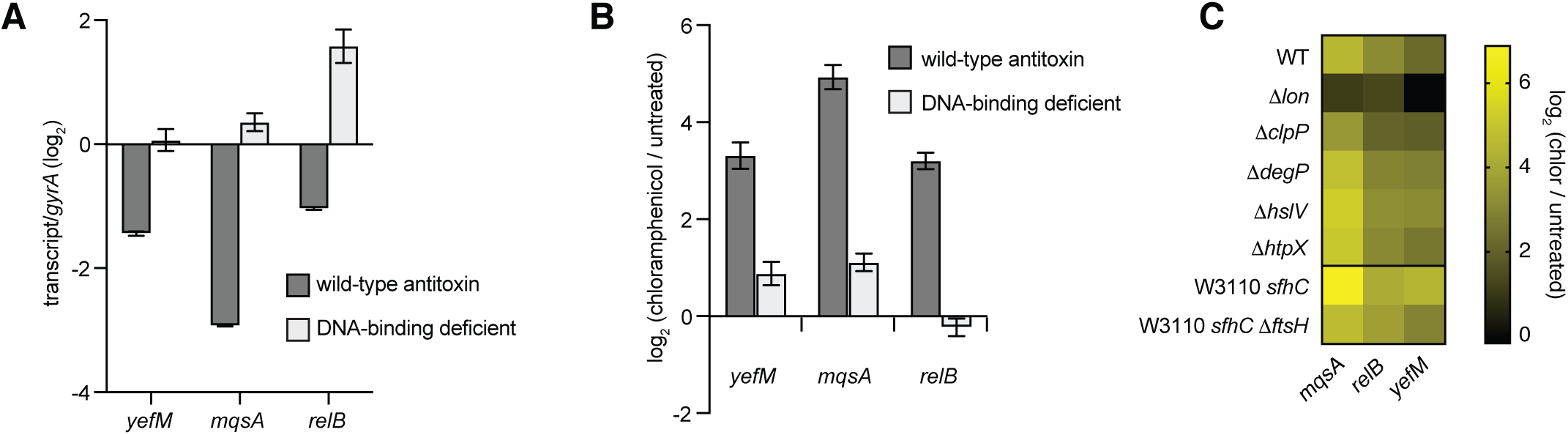
Increases in TA transcription result from a relief of autoregulation. (A) Antitoxin transcript levels were measured in untreated cells at OD_600_ ∼0.3 by qRT-PCR for the wild-type and corresponding DNA-binding mutant allele for the indicated antitoxin. Transcripts were normalized to the *gyrA* housekeeping gene. Data are the average of 3 biological replicates with error bars indicating the S.D. (B) Changes in antitoxin transcript levels in chloramphenicol-treated compared to untreated cells as measured by qRT-PCR. Mutations abrogating DNA-binding by each antitoxin were introduced on the chromosome. Data are the average of 3 biological replicates with error bars indicating the S.D. (C) Changes in antitoxin levels for chloramphenicol-treated cells compared with untreated cells of the wild-type strain or the indicated protease deletion mutant. Transcript levels were quantified by qRT-PCR. Data are the average of 2 biological replicates with error bars representing the S.D.

To investigate whether antitoxin degradation was required for the stress-dependent increases in TA transcription, we screened a panel of *E. coli* protease deletion strains for changes in antitoxin transcript levels following chloramphenicol treatment. Although Lon is often involved in degrading antitoxins, other proteases have been implicated (Aizenman et al., 1996; Christensen et al., 2001; Janssen et al., 2015; Wang et al., 2011). Because an *ftsH* deletion results in a lethal dysregulation of lipopolysaccharide in *E. coli* it can only be generated in a strain carrying a suppressor mutation, *sfhC*; we thus compared the W3110 *sfhC* Δ*ftsH* strain to the W3110 *sfhC* background (Fig. 4C, bottom two rows) (Ogura et al., 1999). Strikingly, the induction of antitoxin transcription by chloramphenicol was largely abolished in a *lon* deletion strain for each of the three TA systems examined (Fig. 4C, Fig. S4). A small reduction in transcription was also evident in the *clpXP* and *ftsH* deletion strains, indicating that these proteases may contribute to the degradation of these antitoxins (Fig. 4C, Fig. S4). Collectively, our results indicate that the increase in TA transcription following stress arises from antitoxin degradation, primarily by Lon, and a consequent relief of autoregulation.

### Transcriptional induction of TA systems does not require the stringent response or ppGpp

A previous model posited that Lon is stimulated to degrade antitoxins at a higher rate following the accumulation of ppGpp (Fig. 5A) (Maisonneuve and Gerdes, 2014). This model claimed that ppGpp promoted the accumulation of polyphosphate, which then directly stimulated Lon to degrade antitoxins, thereby liberating toxins to promote persistence. While results related to persistence were cited in a retraction of this paper, the data related to Lon activation were not and recent literature continues to invoke a connection between ppGpp and TA systems (Christensen-Dalsgaard et al., 2010; Muthuramalingam et al., 2016; Ronneau and Helaine, 2019). Thus, we wanted to test whether the stringent response and ppGpp are required for the transcriptional activation of TA systems.

**Figure 5.**
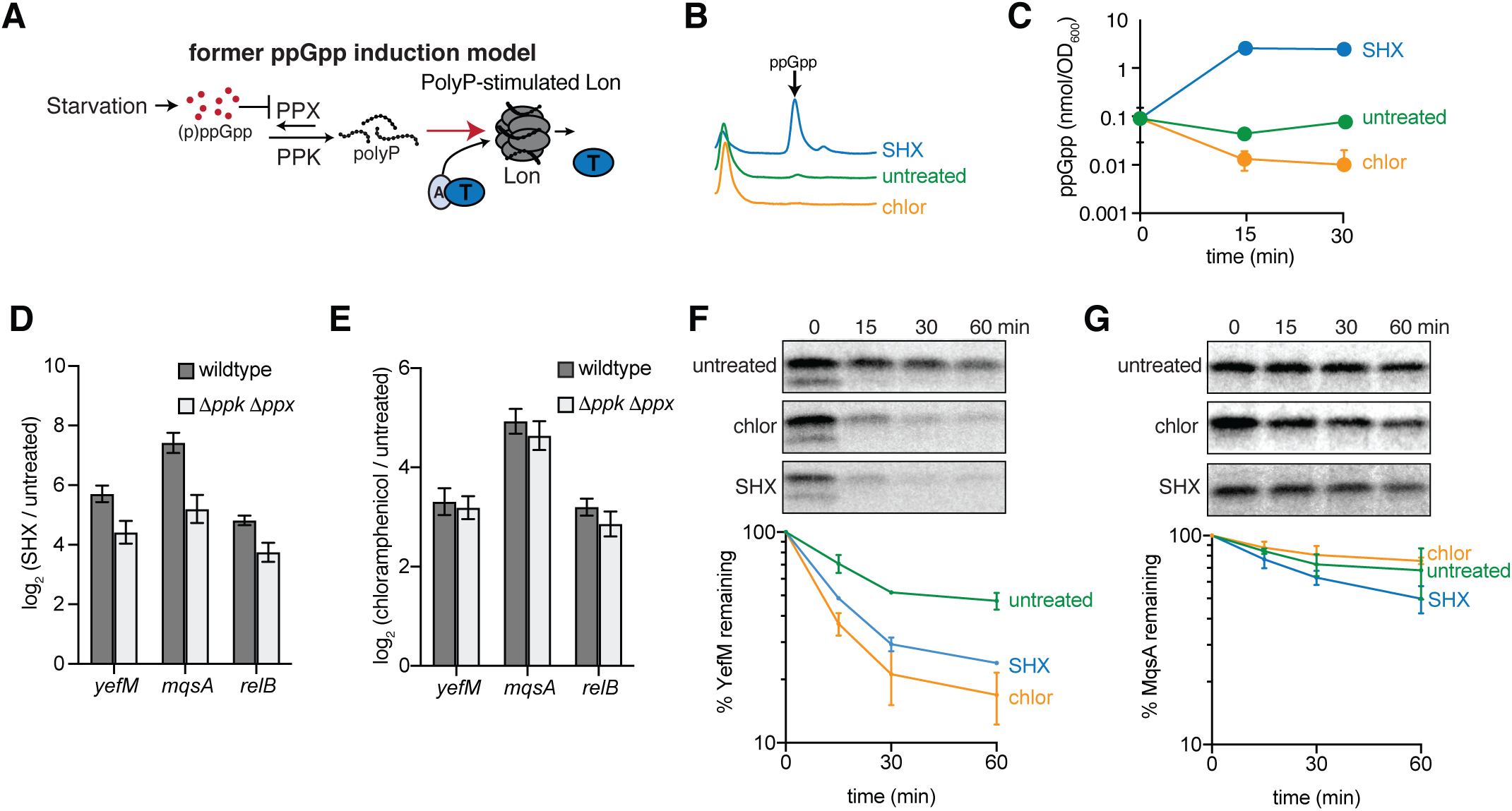
ppGpp is not required for TA transcription. (A) Previous model for ppGpp-mediated activation of TA systems proposed by Maisonneuve et. al. (2013). Amino-acid starvation induces ppGpp, which inhibits polyphosphatase (Ppx), allowing polyphosphate kinase (Ppk) to synthesize polyphosphate, which then stimulates the Lon protease to degrade antitoxins and liberate toxins. (B) Anion exchange chromatography traces for nucleotide extracts from cells harvested 30 min after the indicated treatment. Arrow indicates peak corresponding to ppGpp. (C) Quantification of ppGpp levels in cells after the indicated treatments. Data points are the average of 3 biological replicates with error bars indicating S.D. (D-E) Change in antitoxin transcript levels in cells treated with either SHX (D) or chloramphenicol (E) for 30 minutes compared to untreated cells. Transcript levels were quantified by qRT-PCR. Errors bars represent S.D. and data are the average of 3 biological replicates. (F-G) Antitoxin degradation as measured by pulse-chase analysis. Strains bearing a medium-copy plasmid harboring a TA system under the control of its native promoter, pBR322-*mqsRA* (F) or pBR322-*his_6_-yefM-yoeB* (G), were grown to OD_600_ ∼ 0.3, then pulsed for 10 min with radiolabeled S^35^ cysteine and methionine, and chloramphenicol or SHX added at the same time as the unlabeled cysteine and methionine used to chase. Cell pellets were subsequently lysed and antitoxin was immunoprecipitated and analyzed by autoradiography. Representative gels are shown above with a quantification of at least 2 biological replicates shown below. Error bars indicate S.D.

We showed in Figure 1 that treating cells with SHX, which stimulates ppGpp synthesis, produces a strong transcriptional response. However, treating cells with the antibiotic chloramphenicol, which leads to an accumulation of charged tRNAs that should inhibit RelA to reduce cellular ppGpp levels (Kaplan et al., 1973; J. Sokawa and Y. Sokawa, 1978), also stimulates TA transcription. To determine the levels of ppGpp in our experimental conditions, we treated cells with either SHX or chloramphenicol as before (Fig. 1), and then directly measured ppGpp levels using anion exchange chromatography. After 15 min of SHX treatment, ppGpp levels had reached approximately 2.5 nmol/OD_600_, approximately 27-fold higher than in untreated cells, whereas chloramphenicol treatment reduced ppGpp levels to less than 0.01 nmol/OD_600_, approaching the limit of detection (Fig. 5B-C). We then asked whether cells that had lost the ability to produce polyphosphate – which was proposed to directly activate Lon (Fig. 5A) – could still respond transcriptionally to stress. To test this idea, we used a strain harboring deletions of the genes polyphosphate kinase (*ppk*) and polyphosphate phosphatase (*ppx*), which encode the enzymes responsible for producing and degrading polyphosphate. We found that when this strain was treated with either SHX or chloramphenicol, TA transcription was still induced (Fig. 5D-E). The magnitudes of induction were slightly less than in a wild-type strain, possibly because Δ*ppk* Δ*ppx* cells respond differently to high levels of ppGpp. Nevertheless, this strain exhibited substantial increases in TA transcription following translational stress, regardless of whether ppGpp is produced. Thus, our results indicate that neither ppGpp nor polyphosphate is required for the transcriptional induction of TA systems.

### Translation inhibition can accelerate antitoxin degradation

To more directly test whether ppGpp affects the stability of antitoxins, as postulated previously (Fig. 5A), we used pulse-chase assays to measure the degradation rates of the antitoxins YefM and MqsA in cells experiencing high or low ppGpp levels, induced by treatment with SHX and chloramphenicol, respectively. Native antitoxin levels were too low to be detected by pulse-chase, so we individually cloned the *yefM-yoeB* and *mqsRA* operons – including the native promoter in each case – onto a medium-copy plasmid. There was no substantial increase in the rate of degradation for YefM or MqsA in SHX compared to chloramphenicol (Fig. 5F-G). In fact, MqsA was degraded slightly faster in the low ppGpp (chloramphenicol) condition than in the high ppGpp (SHX) condition (Fig. 5G). These results directly contradict those reported previously (Maisonneuve and Gerdes, 2014), and further indicate that the degradation of antitoxins following stress does not depend on ppGpp.

Our results indicate that TA transcription following chloramphenicol or SHX treatment likely arises from (i) an inhibition of protein synthesis by these stressors and (ii) antitoxin degradation by Lon, leading to a relief of TA autoregulation. In some cases, these stresses accelerate degradation beyond basal, pre-stress rates, though in a ppGpp-independent manner (Fig. 5F-G). Notably, many prior studies of antitoxin stability have relied on Western blotting of chloramphenicol-treated cells, so-called translational shut-off assays. Our finding that chloramphenicol-treatment significantly accelerates the degradation of YefM (Fig. 5F) and moderately increases the degradation of MqsA (Fig. 5G) indicate that such shutoff assays may confound the accurate assessment of degradation rates.

### Heat shock induces transcription of TA systems by increasing antitoxin degradation

In contrast to chloramphenicol and SHX treatment, some stresses that lead to TA transcription, *e.g.* heat shock (Fig. 1), do not inhibit protein synthesis. In these cases, the stress presumably must accelerate the degradation of antitoxins to produce the transcriptional induction observed. To test this hypothesis, we used pulse-chase analyses to examine the effect of heat shock on antitoxin stability following a shift from 30 to 45 °C. Indeed, we found that heat shock significantly increased the rate of YefM degradation (Fig. 6A, S5A), consistent with prior studies (Janssen et al., 2015). Because heat shock does not inhibit translation, the increased degradation of YefM and consequent relief of autorepression increases antitoxin transcription (Fig. 1) and should also lead to an increase in antitoxin protein levels. To test this prediction, we examined YefM levels following heat shock. We were unable to detect YefM expressed from its native promoter by Western blot, so we radiolabeled proteins to steady state and measured YefM levels by immunoprecipitation and phosphorimaging. Following heat shock, the levels of YefM increased approximately 2-fold over 60 minutes, as predicted (Fig. 6B, S5B).

**Figure 6.**
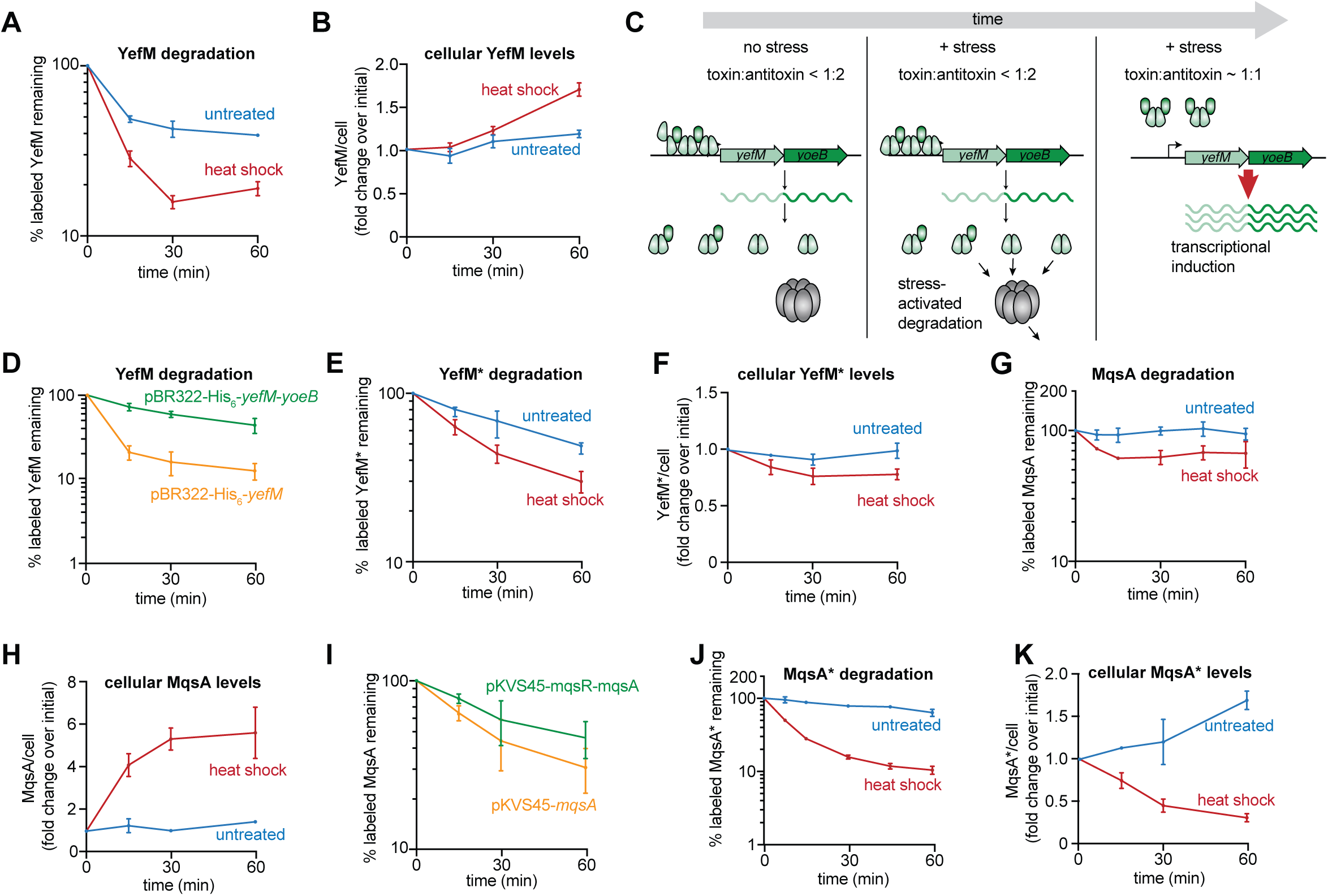
Autoregulation maintains antitoxin homeostasis despite changes in degradation. (A) Antitoxin degradation rate of His6-YefM (A) measured by pulse-chase analysis in cells grown at 30 °C and then either kept at 30 °C (untreated) or shifted to 45 °C (heat shock). (B) Cellular antitoxin levels were measured for His6-YefM by steady-state radiolabeling followed by immunoprecipitation and autoradiography using the same growth conditions as in panels A. (C) Model for transcriptional induction of yefM-yoeB following a stress like heat shock. Stress leads to increased degradation of YefM antitoxin, particularly free antitoxin. As the ratio of toxin:antitoxin increases, YefM antitoxin cannot efficiently bind the yefM-yoeB promoter, leading to sustained transcriptional induction. (D) Degradation rates measured by pulse-chase analysis for His6-YefM expressed with its cognate toxin, YoeB (green) or by itself (orange). (E) Antitoxin degradation rate of His6-YefM* measured as described in panel A. (F) Cellular antitoxin levels of His6-YefM* measured as described in panel B. (G) Antitoxin degradation rate of MqsA measured as described in panel A. (H) Cellular antitoxin levels of MqsA measured from wild-type cells by western blot. (I) Degradation rates measured by pulse-chase analysis for MqsA expressed with its cognate toxin, MqsR (green) or by itself (orange). (J) Antitoxin degradation rate of MqsA* measured as described in panel A. (K) Cellular antitoxin levels of MqsA* measured as described in panel H.See also Fig. S4.

Notably, wild-type cells sustain transcriptional induction (Fig. 1) and continue to accumulate antitoxin after stress (Fig. 6B), indicating that the newly synthesized antitoxin does not immediately feedback and inhibit transcription. This sustained induction likely results from the conditional cooperativity property of many TA systems including YefM-YoeB (Fig. 6C). *In vitro* studies of YefM-YoeB found that when toxin:antitoxin ratios are ∼1:2, the toxin promotes binding of antitoxin to its own promoter; however, at a toxin:antitoxin ratio > 2:1, the toxin disrupts DNA binding by the antitoxin (Kedzierska et al., 2006). Thus, we hypothesized that an increase in the toxin:antitoxin ratio occurs after heat shock, producing the sustained transcriptional induction observed (Fig. 6C). Our attempts to directly measure the toxin:antitoxin ratio were unsuccessful as adding an epitope tag to the toxin affected the toxin:antitoxin interaction. We therefore took a different approach to testing our hypothesis. We reasoned that an increase in the toxin:antitoxin ratio would occur only if free antitoxin were preferentially degraded compared to antitoxin in complex with toxin. Indeed, we noticed that although SHX, chloramphenicol, and heat shock all increased the degradation rate of YefM, ∼20% of the initially labeled pool remained stable for up to 60 minutes (Fig. 5F, Fig. 6A). We suspected that this stable pool is YefM antitoxin bound to YoeB toxin.

To test whether YoeB affects YefM stability, we used pulse-chase analysis to measure the stability of YefM expressed alone from a plasmid, or co-expressed with its cognate toxin YoeB, in the absence of any stress. We found that YefM was significantly less stable in the absence of YoeB, with labeled YefM nearly undetectable after 15 minutes (Fig. 6D, Fig. S5C). The preferential degradation of free antitoxin supports a model in which the ratio of toxin to antitoxin increases following a stress that promotes antitoxin degradation. Additionally, the relative stability of toxin:antitoxin complexes ensures that little to no free toxin is released. Taken together, these results indicate that although stress can trigger a significant and sustained transcriptional induction of *yefM-yoeB*, it does not result in the liberation of significant levels of free toxin (Fig. 6C).

Finally, because the increase in YefM levels following heat shock results from a relief of autoregulation, it should depend on the DNA-binding ability of YefM. To test this idea, we examined strains expressing the DNA-binding deficient YefM variant, YefM*. We first measured changes in YefM* degradation rates in response to heat shock, and found that, as with the wild-type protein, heat shock accelerated the degradation of YefM* compared to untreated cells (Fig. 6E, Fig. S5D). We then measured overall protein levels of YefM* by immunoprecipitation and found that the levels did not increase as with wild-type YefM, but instead decreased to about 80% that seen in untreated cells (Fig. 6F, Fig. S5E). This result confirms that heat shock in wild-type cells triggers accelerated degradation of YefM antitoxin, leading to a relief of transcriptional autoregulation.

We also tested how heat shock affects the MqsA-MqsR TA system. As with YefM, we observed an increase in MqsA antitoxin degradation following heat shock (Fig. 6G, Fig. S5F). When we examined MqsA levels by Western blot, we detected a six-fold increase in MqsA under heat shock conditions compared with untreated cells (Fig. 6H, Fig. S5G). The MqsA-MqsR system does not exhibit conditional cooperativity. However, like the YefM-YoeB system, as the concentration of MqsR toxin increases, it disrupts the binding of MqsA to DNA (Brown et al., 2013). Thus, the sustained induction of *mqsRA* following stress likely arises, as with YefM-YoeB, from increased degradation of MqsA leading to increases in the ratio of MqsR:MqsA, with MqsA bound to MqsR unable to efficiently autoregulate transcription (Fig. S5H). We noted that MqsA degradation also plateaued like YefM following heat shock, leveling off at approximately 60% initial levels, suggesting that a pool of MqsA bound to MqsR may be recalcitrant to degradation. Consistent with this idea, MqsA is degraded more rapidly when expressed alone rather than with its cognate toxin, MqsR (Fig. 6I, Fig. S5I). As with YefM, the preferential degradation of free MqsA antitoxin relative to MqsA bound to MqsR likely helps ensure that no toxin is liberated following stress (Fig S5H).

We also measured the degradation and cellular abundance of MqsA* following heat shock. As with YefM*, MqsA* was degraded more rapidly after heat shock, and cellular levels decreased approximately 2-fold over the course of 60 minutes, indicating that cellular increases in MqsA abundance result from a relief of autoregulation (Fig. 6K, Fig. S5K). We also found that MqsA* is significantly more labile than wild-type MqsA both at 30 °C and following heat shock, suggesting that DNA binding may normally help protect MqsA from degradation, as reported for other Lon substrates (Pruteanu et al., 2007; Shah and Wolf, 2006).

### Chromosomal TA systems differ from plasmid-based systems

Our results suggest that the *E. coli* chromosomal TA systems are fundamentally different from plasmid-based systems. For plasmid-based systems such as CcdAB, toxins are presumably liberated when synthesis abruptly stops following plasmid loss (Van Melderen et al., 1994) as plasmid-free cells are inviable. Our results suggest that a similar mechanism would not be sufficient to activate chromosomal TA systems, as these antitoxin-toxin complexes are relatively resistant to proteolysis, preventing toxin from being liberated. To directly compare the chromosomal systems to CcdAB in a scenario similar to plasmid loss, we generated strains in which the only copy of MqsRA, YefM-YoeB, or CcdAB was under control of the arabinose promoter, P*_ara_*, on the chromosome. Each strain was either grown in the presence of arabinose to express the TA system or not induced. We then rapidly shut off new synthesis (analogous to plasmid loss) by washing cells and releasing them into medium containing glucose, which represses P*_ara_* (Fig. 7A). As expected, cells harboring *ccdAB* showed a clear decrease in growth rate ∼4 hrs after inhibiting synthesis compared to cells in which CcdAB was not initially induced (Fig. 7B). The slow time scale of CcdAB activation is consistent with early studies of plasmid loss (Jaffé et al., 1985), and likely represents a combination of time required for antitoxin turnover and the time needed for cells to adapt to a change in carbon source (from glycerol + arabinose to glucose). In contrast to CcdAB, we saw no changes in growth following the induction and subsequent repression of either MqsRA or YefM-YoeB (Fig. 7C-D). These findings are consistent with our pulse-chase studies (Fig. 5-6) indicating that the antitoxins MqsR and YefM are, in contrast to CcdA, maintained in relatively stable complexes with their toxins such that blocking new synthesis does not lead to their rapid decay and a consequent release of growth-inhibiting toxin.

**Figure 7.**
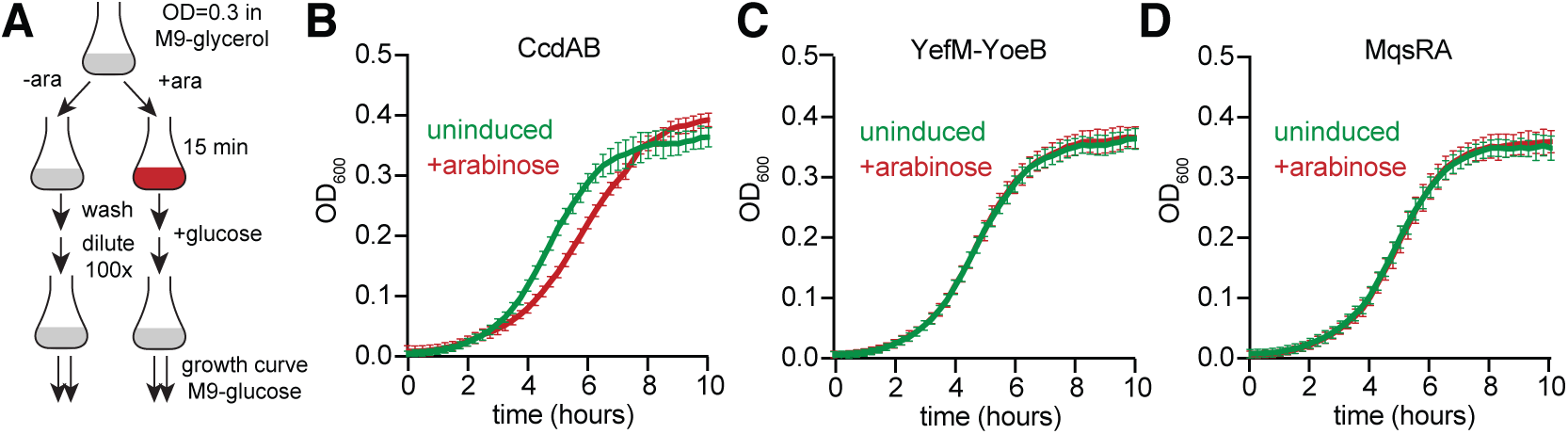
Chromosomal TA systems are not induced by an inhibition of new synthesis as with the plasmid-borne system CcdAB. (A) Schematic for growth experiments in panels B-D. Strains were grown to OD_600_ ∼ 0.3 in flasks, then split into 2 flasks, one of which was induced with 0.2% arabinose for 15 min. After washing, cells were released into repressing conditions (M9 medium containing glucose) at a 100x dilution and growth was measured on a plate reader. (B-D) Growth curves resulting from experiment described in panel A for (B) *attL*::*P_ara_-ccdAB*, (C) Δ*yefM-yoeB attL*::*P_ara_-yefM-yoeB*, and (D) Δ*mqsRA attL*::*P_ara_-mqsRA*.

## Discussion

Contrary to the widely-held view that TA systems function as stress-response modules, we found that abiotic stress can trigger transcription of chromosomally encoded TA systems in *E. coli*, but that significant, growth-inhibitory levels of free, active toxin are not generated by these stresses. Numerous prior studies have interpreted the robust increases in TA transcription to mean that (i) antitoxin levels were depleted and (ii) that this results in release of active toxin (Christensen-Dalsgaard et al., 2010; Muthuramalingam et al., 2016; Ronneau and Helaine, 2019). Our results support the initial inference that stress can accelerate antitoxin turnover and thereby induce transcription by relieving autorepression. However, we find that, despite increases in antitoxin degradation rates, antitoxin pools are never fully depleted, and thus toxin is never actually freed. Using two different measures of toxin activity: changes in growth, and mRNA cleavage, we find no evidence that the 10 endoribonuclease toxins of *E. coli* are significantly activated by numerous, diverse abiotic stresses on time-scales coincident with major changes in their transcription.

How can an increase in antitoxin turnover occur, yet not liberate toxin? Two major factors contribute: (i) antitoxins are typically much more abundant than their cognate toxins and (ii) free antitoxin is preferentially degraded relative to antitoxin bound to its cognate toxin. Thus, in cases where a stress, e.g. heat shock, triggers accelerated degradation of antitoxin, the ratio of toxin:antitoxin increases. For both YefM-YoeB and MqsA-MqsR, increases in this ratio are known to disrupt autorepression by the antitoxin (Brown et al., 2013; Kedzierska et al., 2006), thereby explaining the sustained transcriptional induction. Importantly, because antitoxin bound to toxin is recalcitrant to degradation, there is little to no free toxin released, despite the transcriptional induction. In cases where a stress does not accelerate degradation but also blocks new synthesis, e.g. chloramphenicol treatment, the same logic applies. Free antitoxin is preferentially degraded, leading to an increase in the ratio of toxin:antitoxin, which will drive transcriptional induction, but without commensurate increases in protein levels given the block to synthesis. In sum, our results now provide a mechanistic basis for understanding how diverse stresses can trigger transcriptional induction, and in some cases increased protein levels, without liberating active toxin.

### Stress-induced changes in antitoxin degradation

How is Lon-dependent degradation of antitoxins stimulated by diverse stresses? Early studies of Lon suggested that it can be activated by polyphosphate to degrade ribosomal proteins (Kuroda et al., 2001). A subsequent study extended this idea, postulating that the ppGpp made during amino acid starvation can stimulate polyphosphate to activate Lon-dependent degradation of antitoxins (Maisonneuve et al., 2013). However, we found no evidence that either ppGpp or polyphosphate are required for the increased degradation of antitoxins in stressful growth conditions. An alternative possibility is that because Lon normally degrades misfolded proteins that arise during translation (Mahmoud and Chien, 2018; Van Melderen and Aertsen, 2009), whenever the abundance of misfolded or incompletely synthesized peptides is substantially reduced, Lon may be more accessible to lower affinity substrates, such as antitoxins. The relatively long half-life of many antitoxins (>20 min) compared to some other Lon substrates (e.g. SulA, ∼1 min) (Mizusawa and Gottesman, 1983), suggests that they may indeed be lower affinity Lon substrates, and it is known that Lon saturation affects the rate at which it degrades lower affinity substrates (Dervyn et al., 1990). Most of the stresses we tested were severe enough that protein translation was likely either fully or strongly inhibited. Exceptions to this include heat shock and the deletion of *dnaK*, although both of these conditions, which lead to proteotoxic stress, are known to stimulate Lon (Janssen et al., 2015; Jonas et al., 2013). A second possibility is that a Lon activator could become more abundant during stress. A subset of Lon substrates can function as activators of Lon in *Yersinia pestis* and *Caulobacter crescentus* (Jonas et al., 2013; Puri and Karzai, 2017). Because we saw an increase in antitoxin degradation even when translation is fully inhibited, this is a less likely scenario. However, we cannot rule out that a Lon activator is stabilized or becomes more abundant by a synthesis-independent mechanism.

Our studies of antitoxin degradation have also uncovered two important technical considerations when measuring protein half-life. First, the fact that antitoxins can be degraded faster, as measured by pulse-chase analysis, when cells are exposed to translation-inhibiting antibiotics is of broad relevance, given the common practice of measuring protein turnover by Western blotting after treating cells with antibiotics such as chloramphenicol to shutoff new translation. Because Lon activity increases in response to such antibiotics, this method may significantly overestimate the lability of some Lon substrates. A second important finding is that the half-life of an antitoxin depends on how it is produced. For instance, we found that the half-life of an antitoxin can depend strongly on whether its cognate toxin is being co-expressed or not (Fig. 6J). Similarly, we found that interactions with DNA can significantly affect the half-life of antitoxins, with point mutations that disrupt DNA-binding by antitoxins leading to increased lability (Fig. 6G and 6J). Similar effects have been documented for other Lon substrates (Ahn and Baker, 2016; Pruteanu et al., 2007; Shah and Wolf, 2006).

### Functions of toxin-antitoxin systems

Our results demonstrate that diverse stresses can trigger increased degradation of antitoxins and that this relieves autoregulation to drive transcriptional induction of TA operons. However, these dynamics do not lead to the liberation of significant amounts of active toxin. If free toxin were being produced at appreciable levels, we would have seen a TA-dependent suppression of, or delay in, growth following the application of a stress. However, we observed no detectable reduction in the growth of wild-type cells compared to cells lacking 10 TA systems following heat shock, acid shock, or treatment with chloramphenicol, hydrogen peroxide, trimethoprim, or SHX (Fig. 2). As the 10 TA systems deleted each feature an endoribonuclease toxin, we also used RNA sequencing to test whether there is any evidence of toxin activity, but no cleavage of mRNA was detected (Fig. 3). It could be that TA systems are activated only in a very tiny fraction of cells experiencing a given stress such that no changes in bulk growth or RNA cleavage can be detected. But such an argument would still mean that TA systems are not being used as general stress-response modules.

How, then, do toxins ever get activated? Presumably, antitoxins must be dissociated (with or without degradation) from their cognate toxins, but the conditions that trigger such dissociation and, consequently, activation of a toxin remain unclear for most chromosomally-encoded TA systems. We find that the chromosomal type II TA systems function differently from the well-characterized plasmid-borne TA system CcdAB that inhibits cell growth upon plasmid loss (Van Melderen et al., 1994). The cessation of *ccdAB* expression following plasmid loss, coupled with the relative instability of CcdA, leads to the liberation of CcdB toxin, an effect that can be recapitulated by shutting off synthesis of *ccdAB* (Fig. 7B). In contrast, we found that inhibiting synthesis of the chromosomally-encoded *mqsRA* and *yefM-yoeB* systems was not sufficient to activate toxin, as measured by cell growth.

It remains to be seen whether plasmid-borne systems have different roles from chromosomal systems, or whether chromosomal systems are plasmid-derived systems that have lost, or are in the process of losing, their function over evolutionary time. We think the latter is unlikely because all the *E. coli* toxins have retained full activity. Another possibility is that the chromosomal systems in *E. coli* are activated by the same mechanism as CcdAB, but that they have slower off rates, and therefore require considerably longer timescales for toxin to be freed. One result in line with this model is the observation that the DinJ-YafQ system may protect cells against cefazolin or tobramycin when they are grown as biofilms, but not when grown in liquid (Harrison et al., 2009). Biofilms take several days to develop, and cells at the interior of the biofilm experience nutrient-poor conditions that may be sufficiently stressful to stimulate antitoxin degradation and prevent synthesis of new antitoxin over extended periods of time.

It is also conceivable that additional signals are needed to destabilize chromosomally-encoded TA complexes. Such a signal might arise under conditions not tested here, e.g. a biotic stress. A biotic stress could consist of phage infection, bacterial residence within a eukaryotic cell, or interbacterial antagonism within a microbial community. Each of these environments or conditions could lead to the co-occurrence of multiple stresses that induce toxin activity or the production of an unknown signal that alone is sufficient to induce toxins. Notably, the most recently discovered type II TA system in K12 *E. coli,* RnlAB, plays a clear role in defending *E. coli* against T4 phage infection, indicating that this system must somehow become active during phage infection (Koga et al., 2011), but the mechanism responsible is not known. Additionally, the chromosomal type II TA systems in *Salmonella enterica* ser. Typhimurium help promote survival within macrophages, indicating that they are active in this environment (Helaine et al., 2014), but again a mechanism of induction is not known.

In sum, our results emphasize that TA systems are unlikely to be critical or conserved components of bacterial stress response systems. Additionally, our findings strongly underscore the notion that the transcriptional activation of TA systems is not a reliable marker for activity. Specific assays of toxin activity such as the RNA-seq method used here to probe endoribonuclease activity will be crucial for identifying and characterizing the *bona fide* inducers of TA systems.

## Acknowledgements

We thank M. Guzzo, K. Gozzi, S. Jones, and M. Guo for comments on the manuscript and C. Aakre and D. Huang for helpful discussions. M. LeRoux was a Simons Foundation Fellow of the Life Sciences Research Foundation. This work was funded by an NIH grant to M.T.L. (R01GM082899), who is also an Investigator of the Howard Hughes Medical Institute.

## Author Contributions

M.L performed all experiments. P.H.C. helped with RNA-seq experiments and analysis. Y.J.L. helped with qRT-PCR experiments and M. Littlehale helped with growth studies. M.L. and M.T.L designed experiments, analyzed data, prepared figures, and wrote the manuscript.

## Supplemental Figure Legends

**Figure S1.**
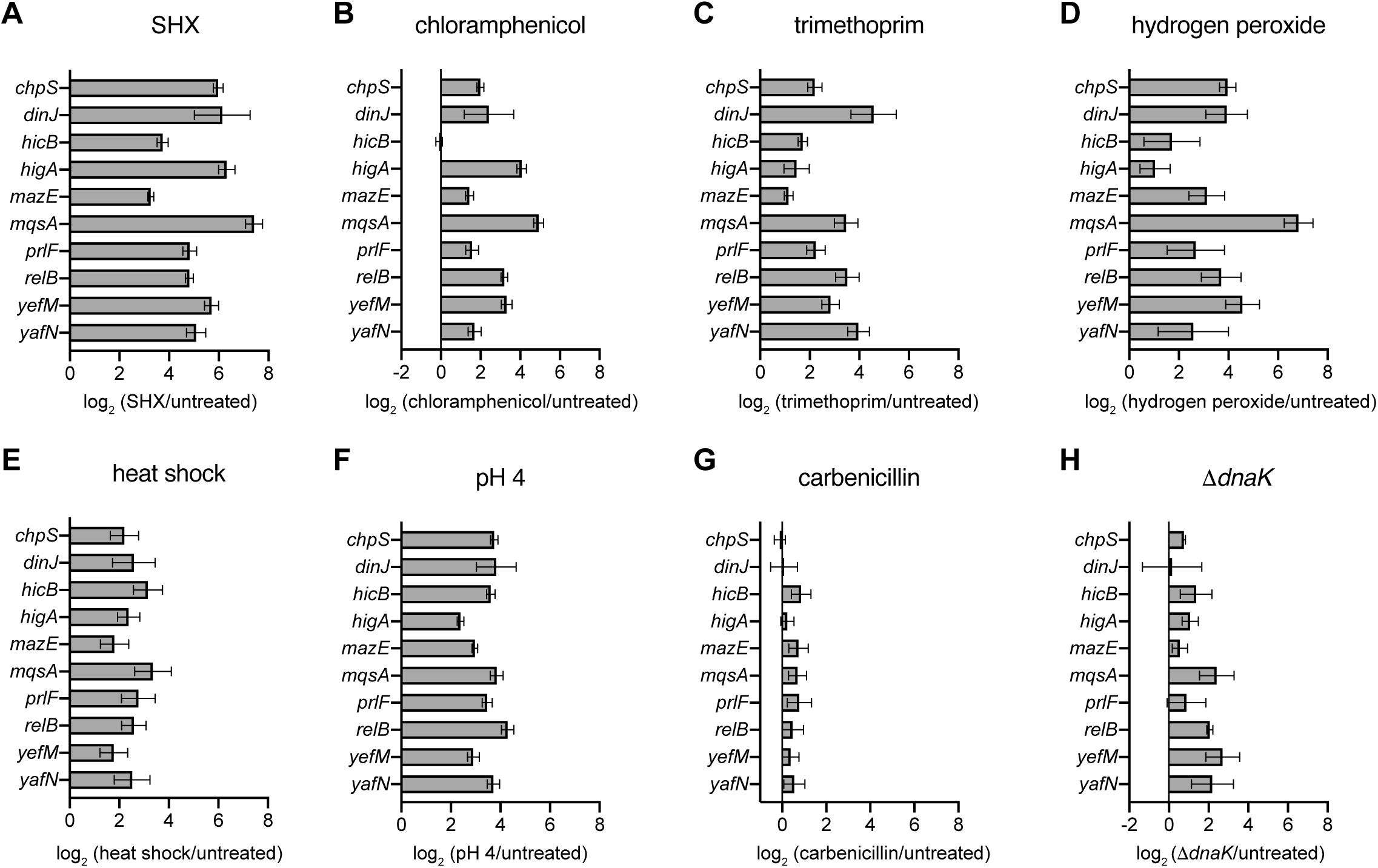
TA systems are transcriptionally activated by diverse stresses. Difference in antitoxin transcript levels between cells exposed to the indicated stress for 30 minutes compared to untreated cells as measured by qRT-PCR corresponding to Figure 1D. Data presented are the average of 3 biological replicates errors bar represent S.D.

**Figure S2.**
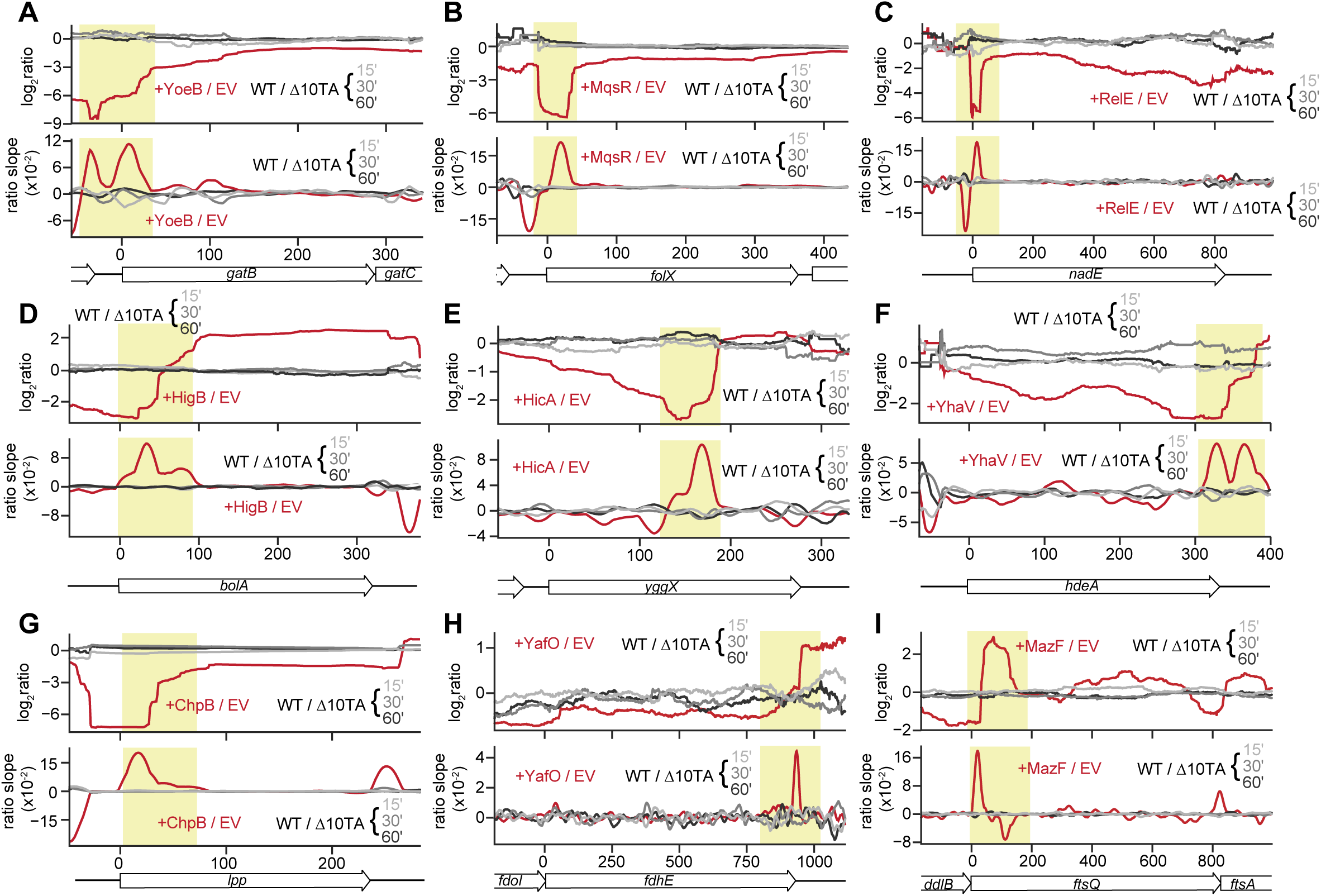
Transcripts targeted by toxins are not cleaved after chloramphenicol treatment. (A-I) Examples of transcripts that show signatures of cleavage following toxin overexpression. Ratios of +/− toxin and chloramphenicol-treated wild type / Δ10TA (upper panel) and corresponding slopes (lower panel) for each of the nine toxins examined. See also Figure 3.

**Figure S3.**
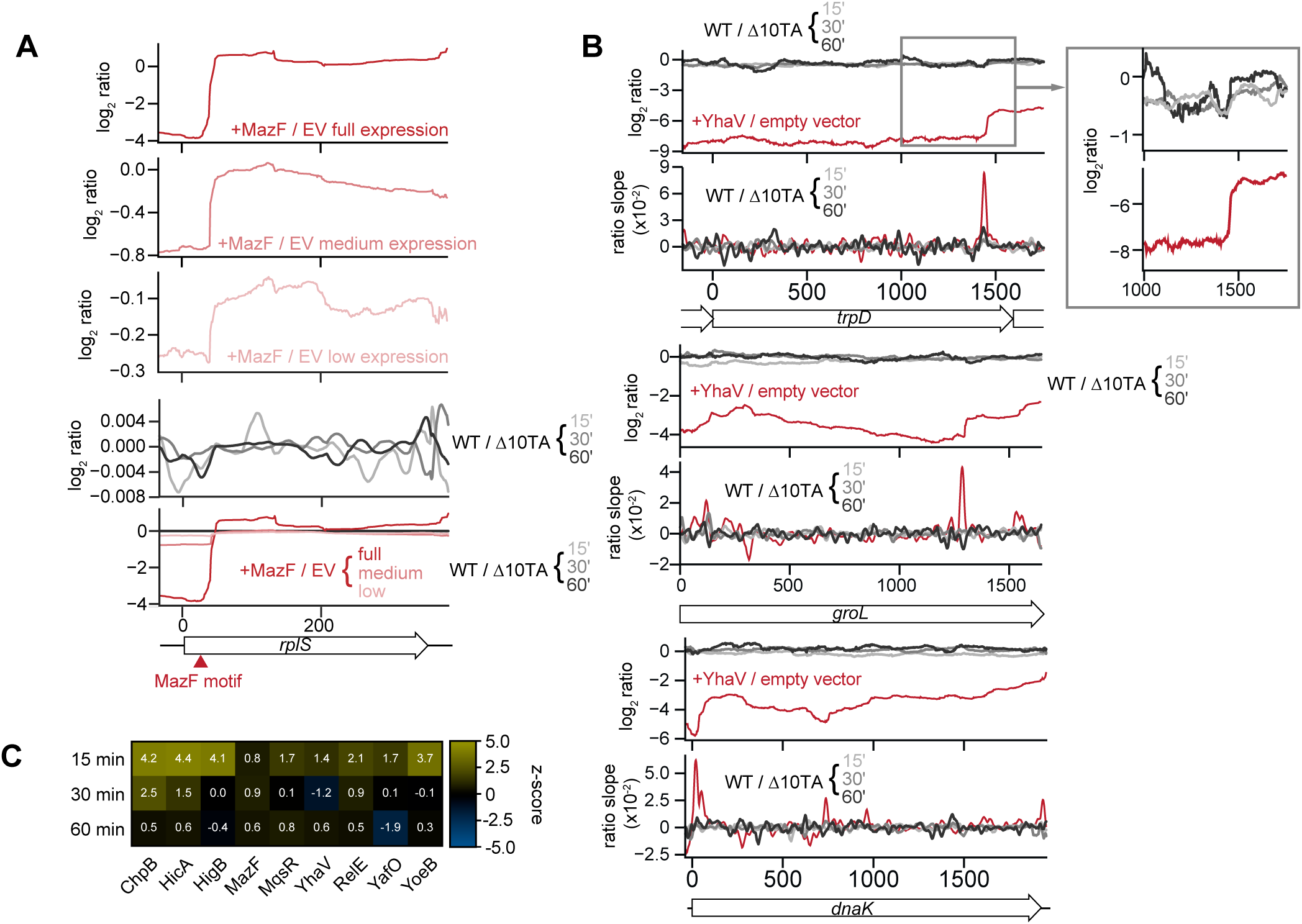
No evidence of toxin cleavage following chloramphenicol treatment. (A) Comparison of of the shape of ratio plots of +MazF / empty vector for *rplS* transcript containing a MazF cleavage site at high, medium, and low MazF expression levels (upper 3 panels) and for wild type / Δ10TA in chloramphenicol treatment samples (fourth panel). Lower panel is an overlay of all five ratio profiles plotted on the same scale. (B) YhaV regions containing the highest cleavage ratio slopes corresponding to analysis described in Fig. 3E. Chloramphenicol-treated samples are compared to YhaV overexpression data. (C) Z-scores generated from comparing the average of the minima detected in the transcripts with the lowest +/− toxin ratios (lowest 5%, from toxin overexpression datasets) to the average of the minima from a randomly selected set of transcripts (sampled 10,000 times) in the chloramphenicol RNA-seq datasets. A negative z-score indicates a significant amount of toxin activity; positive z-scores may result from transcript stabilization. See also Figure 3.

**Figure S4.**
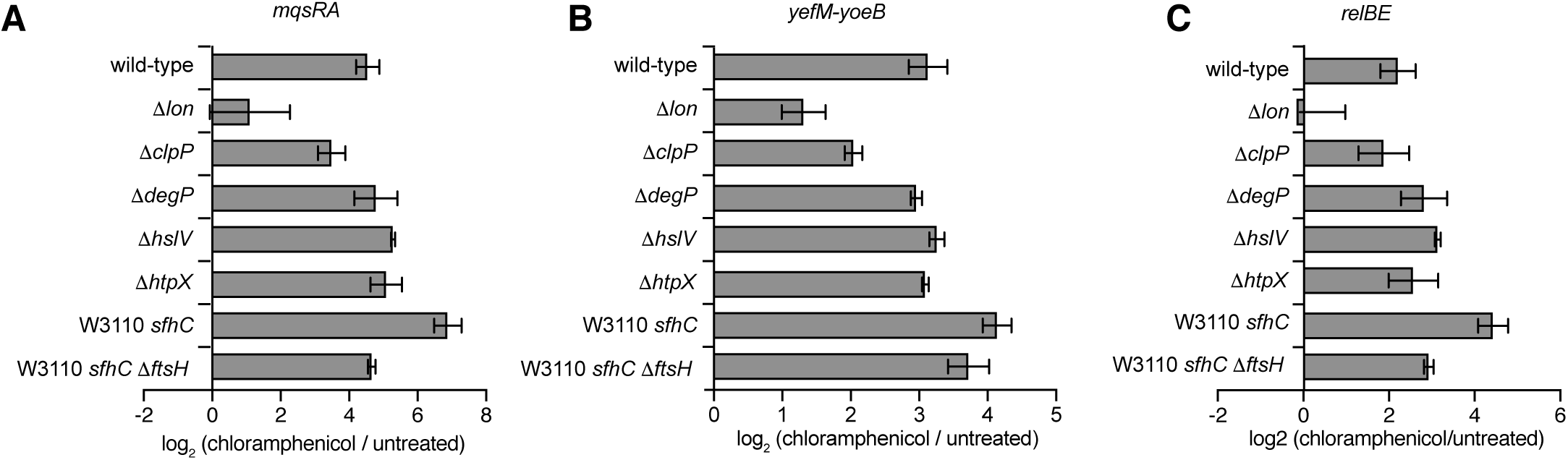
Chloramphenicol-induced changes in transcription require the Lon protease. Difference in antitoxin levels of chloramphenicol-treated compared with untreated cells of the indicated protease deletion mutants for *mqsRA* (A), *yefM-yoeB* (B), and *relBE* (C). Transcript levels were quantified by qRT-PCR and correspond to Fig. 4B. Data presented are the average of 2 biological replicates and error bars represent S.D. See also Figure 4.

**Figure S5.**
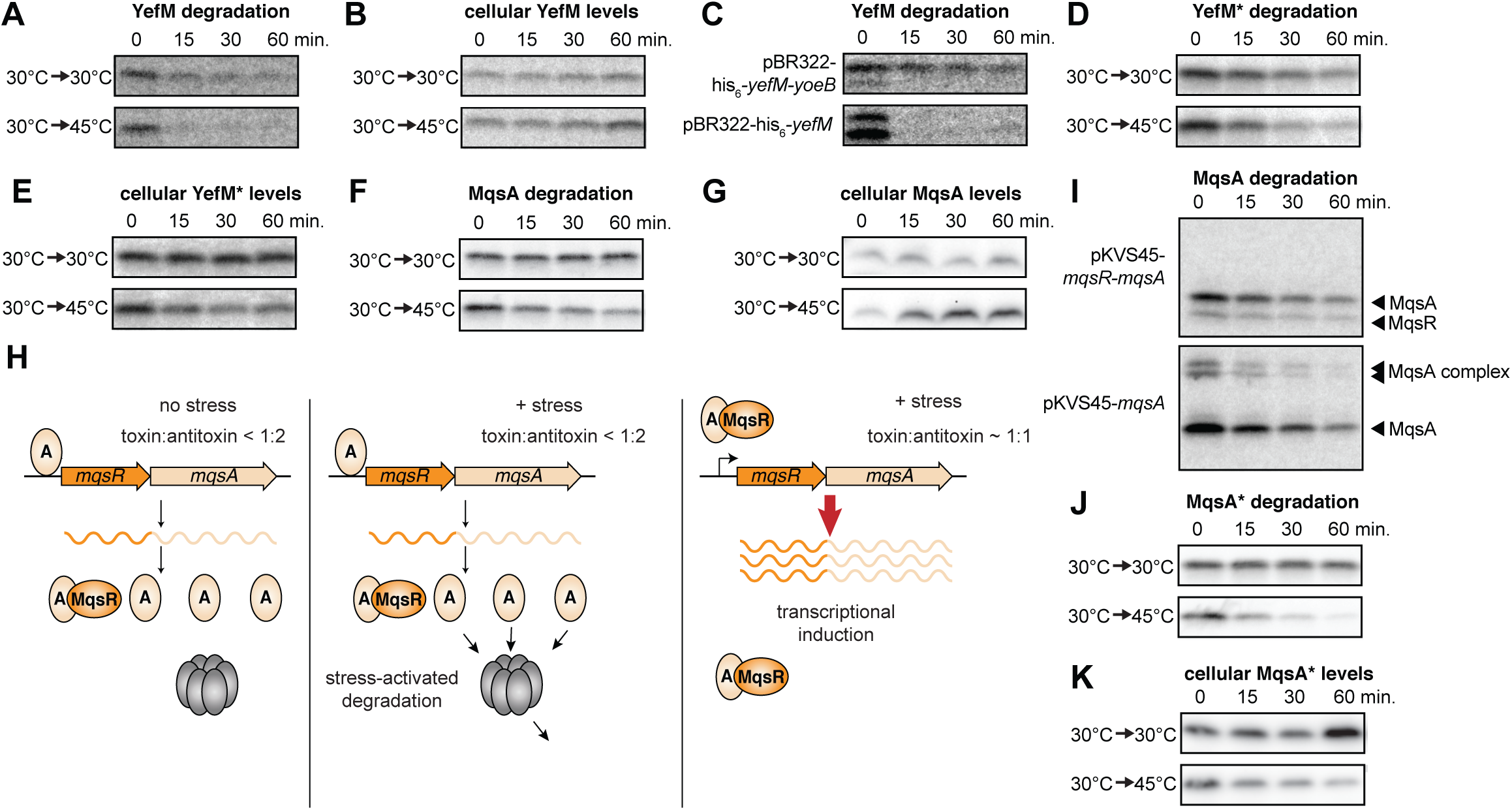
Autoregulation maintains antitoxin homeostasis despite changes in degradation. (A-B) Representative phosphorimages of His_6_-YefM degradation (A) or cellular antitoxin levels (B) in cells grown at 30 °C and then either kept at 30 °C (untreated) or shifted to 45 °C (heat shock). Degradation was measured by pulse-chase and immunoprecipitation while cellular levels were measured by steady-state radiolabeling followed by immunoprecipitation and autoradiography. (C) Representative phosphorimages of His_6_-YefM degradation following co-expression with its cognate toxin, YoeB (upper), or expression by itself (lower). (D-E) Same as panels A-B but for _His6-_YefM*. (F-G) Representative phosphorimages of MqsA degradation measured by pulse chase ((F) or Western blots of cellular MqsA levels (G) under growth conditions described in (A). (H) Model for MqsA protein levels following stress. MqsA cannot bind both DNA and MqsR, therefore at higher degradation rates as toxin:antitoxin levels increase, the promoter is derepressed, leading to transcriptional activation. (I) Representative phosphorimages of MqsA degradation following co-expression with its cognate toxin, MqsR (upper), or expression by itself (lower). (J-K) Same as panels F-G but for MqsA*. See also Figure 6.

## Methods

### Strains and growth conditions

All primers and strains used in this study are listed in Tables S1 and S2, respectively. *E. coli* was cultivated at 37 °C in M9-glucose (6.4 g/L Na_2_HPO_4_-7H_2_O, 1.5 g/L KH_2_PO_4_, 0.25 g/L NaCl, 0.5 g/L NH_4_Cl medium supplemented with 0.1% casamino acids, 0.4% glucose, 2 mM MgSO_4_, and 0.1 mM CaCl_2_) for all experiments, except when otherwise indicated. Routine cultivation for strain construction was in Luria broth (LB). Media for selection or plasmid maintenance were supplemented with carbenicillin (100 µg/mL), chloramphenicol (20 µg/mL), or kanamycin (30 µg/ml) as necessary unless otherwise indicated.

### Plasmid construction

TA systems were inserted by Gibson assembly into pBR322 amplified without the tetracycline promoter using primers ML1 and ML2. The region of the chromosome encoding each TA locus, including approximately 200 bp upstream, was amplified with primers ML3 and ML4 (*yefM-yoeB*), ML5 and ML6 (*mqsRA*), and ML7 and ML8 (*relBE*) with 40 bp ends homologous to pBR322. Mutations previously reported to abrogate DNA binding were introduced into pBR322-*mqsRA*, pBR322-*relBE*, and pBR322-*yefM-yoeB* by site-directed mutagenesis using inverse PCR with primers ML15 and ML16 (*yefM*), ML17 and ML18 (*mqsA*), and ML19 and ML20 (*relB*) resulting in pBR322-*yefM*-yoeB*, pBR322-*mqsRA*(N97A), pBR322-*relB*E*.

pBR322-His_6_-*yefM*-*yoeB* and pBR322-His_6_-*yefM*\*-*yoeB* were constructed by site-directed mutagenesis with inverse PCR using primers ML23 and ML24 using pBR322-*yefM*-*yoeB* and pBR322-*yefM*\*-*yoeB* as templates, respectively. pBR322-His_6_-*yefM* was constructed by inverse PCR using primers ML25 and ML26 with pBR322-His_6_-*yefM*-*yoeB* as the template. pKVS45-*mqsRA* and pKVS45-*mqsA* were constructed using restriction based cloning. MqsRA or MqsA were amplified from MG1655 using primers ML27 and ML28 or ML28 and ML29, respectively, digested with SacI and HindIII, and ligated to pKVS45 digested with the same enzymes.

### Strain construction

Fluorescently labeled wild-type and Δ10TA strains were generated by transduction of *galK::P_lac_-cfp* and *galK::P_lac_-yfp* (Elowitz et. al. 2002). The DNA-binding mutants were constructed by allelic exchange, wherein the TA locus was first replaced with a *sacB-neoR* cassette from pIB279 amplified with primer sets ML9 and ML10 (*yefM-yoeB*), ML11 and ML12 (*mqsRA*), and ML13 and ML14 (*relBE*) (Blomfield et al., 1991). The DNA-binding mutant alleles were then amplified from pBR322-*relB*E*, pBR322-*mqsRA\**, and pBR322-*yefM*-yoeB* described above with homologous ends to the flanking region. Both the counterselectable cassette and subsequent DNA-binding mutant allele were inserted using lambda Red recombinase, either selected on kanamycin or counterselected on 5% sucrose plates as previously described (Datsenko and Wanner, 2000). Mutations were verified by PCR amplification and sequencing.

To make protease deletions, deletion strains were obtained from the Keio collection and transduced into MG1655 with P1 phage. Kanamycin resistance cassettes were removed using pFLP1 excision as previously described (Datsenko and Wanner, 2000). The Δ*ppk* Δ*ppx* strain was constructed by lambda Red to replace the locus with a kanamycin resistance cassette amplified from pKD4 using primers ML21 and ML22 with homology flanking the *ppk ppx* region. The kanamycin resistance cassette was removed with pFLP1 excision.

Arabinose-inducible TA systems were created by in-frame deletion of the TA systems. The *neoR-sacB* cassette replacement strains described above were transformed with corresponding deletion oligos ML36 (*yefM-yoeB*) and ML37 (*mqsRA*) and deletions were counterselected on sucrose plates. TA open reading frames were then inserted downstream of the arabinose promoter of the pAH150 plasmid (Haldimann and Wanner, 2001) by Gibson assembly. TA systems were amplified with primers ML40 and ML41 (*ccdAB*, amplified from the F plasmid); ML42 and ML42 (*yefM-yoeB*), and ML43 and ML44 (*mqsRA*) and pAH150 linearized with primers ML38 and ML39. CRIM insertion was then performed as previously described using helper plasmid pInt-ts with selection on low kanamycin (6 µg/mL) to prevent multiple insertion events (Haldimann and Wanner, 2001). Single insertions were confirmed by PCR as described.

### Stress conditions

For all stress conditions used in qRT-PCR, pulse-chase, immunoblots, and growth experiments, stresses were applied as follows: chloramphenicol at 75 µg/mL, SHX at 100 µg/mL, trimethoprim at 25 µg/mL, H_2_O_2_ (Sigma) at 0.021% vol/vol (7 mM), carbenicillin at 25 µg/mL. For low pH experiments, cells were washed into M9-glucose adjusted to pH of 4 for the duration of the stress treatment. For all heat shock experiments, cells were grown at 30 °C prior to heat shock, then either kept at 30 °C (untreated) or shifted to 45 °C (heat shock).

### qRT-PCR experiments

Cells were harvested by adding 900 µL bacterial culture to 100 µL stop solution (5% phenol, 95% ethanol v/v) on ice. RNA extractions, reverse transcription, and qRT-PCR were performed as described previously (Culviner and Laub, 2018) with the primers listed in Table S1. Briefly, RNA was reverse-transcribed with first-strand synthesis kit (Invitrogen) using random primers. cDNA was diluted 1:15. Standard curves made by combining and diluting cDNA from each set of cDNAs were run for all primers on each plate. Reactions were prepared using 2X SYBR Fast Mastermix (Roche) and run on a Light Cycler 480 II Real-time PCR Machine at the MIT BioMicro Center in technical duplicates. Samples were analyzed using the standard-curve method and transcripts of interest were normalized to the *gyrA* housekeeping gene. All qRT-PCR data reported are the average of biological triplicates, with the exception of the protease deletion strains, which are the average of biological duplicates.

### Growth experiments

For growth experiments, the wild type used was the parental strain to the Δ10TA strain obtained from L. Van Melderen (Goormaghtigh et al., 2018). Cells were grown to OD_600_ ∼ 0.3 and then stresses were applied as described above. After the indicated stress treatment, cells were washed 2x in an equal volume of media. Cells were diluted to OD_600_ of 0.01 in M9-glucose. Growth was measured in 96-well plates at 15 min intervals (180 µL culture overlaid with 70 µL mineral oil) with orbital shaking at 37 °C on a plate reader (Biotek). Representative data presented are the result of 6-12 plate replicates and were replicated independently at least 2 times. For growth measurements during heat shock, cells were diluted to OD_600_ ∼0.01 and then growth curves were performed on a plate reader set either to 30 °C or 44 °C. For the competition experiments, overnight cultures of fluorescently labeled wild-type and Δ10TA were mixed at a 1:1 ratio, then treated to chloramphenicol as described for growth curve experiments. After the first chloramphenicol treatment, cultures were washed and back-diluted in M9-glucose to an OD_600_ of 0.05 and allowed to reach OD_600_∼0.3 before the second chloramphenicol treatment, second wash, and final growth step. Samples were taken at indicated time points, including after overnight growth. Each sample was serially diluted and plated on non-selective plates. Plates were scanned on a Typhoon imager and CFP vs YFP colonies determined using the CY3 fluorescent channel. The experiment was performed with three biological replicates.

### RNA-seq and analysis

Cells were collected and RNA harvested as described above. RNA-seq libraries were prepared as described previously (Culviner and Laub, 2018). Briefly, rRNA was removed with a RiboZero kit (Illumina) per manufacturer instructions. RNA was fragmented with RNA Fragmentation reagents (Invitrogen). Fragmentation reagent was added, then samples were incubated at 70 °C for 8 min before reactions were stopped. Fragmented RNA was ethanol precipitated, then reverse-transcribed using random primers (Invitrogen First Strand Kit and Random Primers). Second strand synthesis was subsequently performed (Invitrogen Second Strand Synthesis kit) and RNA digested with Rnase H (NEB). Paired-end sequencing was performed on an Illumina NextSeq 500 at the MIT BioMicroCenter. Data were uploaded to NCBI Geo (accession number GSE141320).

### RNA-Seq Analysis

RNA-Seq read mapping and calculation of + toxin : empty vector ratios were conducted as described previously (Culviner and Laub, 2018). Slopes were calculated by a least-squares fit of the 30 nucleotides to the 5’ of a given position. To systematically search for cleavage across the transcriptome, we first identified coding genes that had at least 64 counts at all positions across the coding region in the empty vector sample and at least 1 count in the expressed toxin sample. Toxin-antitoxin genes as well as the phase-variable *flu* gene were excluded from this analysis regardless of their expression level. For slope-based identification of cleavage targets, the maximum slope (calculated as described above) was calculated for each transcript. The transcripts with the top 5% of maximum slopes were classified as toxin targets. To determine if cleavage was occurring in the presence of chloramphenicol, we took the ratio profiles for wild type v. Δ10TA in chloramphenicol and calculated the mean of the maximum slopes for the defined set of toxin targets and compared this value to the mean of the maximum slopes for randomly selected transcripts. To do this, we sampled sets of randomly chosen genes (the same number as the number of toxin targets) 10,000 times and calculated the mean of their maximum slopes. In this way, we generated a distribution of the expected value of mean maximum slopes. These distributions are shown as box and whisker plots displaying the median, lower and upper quartiles, and quartile ± 1.5 interquartile range (Fig. 3D). The mean maximum slope of toxin targets was compared to these distributions to calculate a z-score. The above analysis was also modified to detect cleavage by + toxin : empty vector ratios. In this case, the bottom 20% of transcript minimum ratios were classified as toxin targets. The rest of the analysis was conducted as above, but minimum ratios replaced maximum slopes.

### ppGpp measurements

Cells were grown to OD_600_ of 0.2-0.3, then treated with either chloramphenicol or SHX as described, and harvested and quantified as described previously (Kraemer et al., 2019). Briefly, cells were vacuum filtered onto a nitrocellulose membrane and immediately exposed to cold lysis solvent (40% methanol, 40% acetonitrile, 20% water), removed from the membrane by sonication, and flash frozen. Volumes were adjusted based on OD_600_ of the culture at the time of harvest, and ppGpp was quantified from the resulting nucleotide extracts by anion-exchange chromatography (MonoQ 5/50 column) and concentration calculated based on a standard curve. Data presented are the result of biological triplicates.

### Pulse chase experiments

Cells were grown in M9-glucose to OD_600_ ∼ 0.3. Cultures were labeled with 80 µCi/mL [^35^S] EasyTag™ EXPRESS^35^S Protein Labeling Mix (Perkin Elmer) for 10 min, then chased with 2 mM cysteine and methionine, and the indicated stress applied. For all time points, 1 mL of cells were pelleted and flash frozen in liquid nitrogen. Pellets were resuspended in 500 µL lysis buffer (PBS supplemented with 0.05% Tween-20, 1 µL/mL benzonase, 1 µL/mL ReadyLyse (Novagen), and cOmplete protease inhibitor cocktail (Sigma)). Cells were lysed by additional freeze/thaw cycles, cell debris was removed by centrifugation, and proteins were immunoprecipitated with Protein A Dynabeads (ThermoFisher) conjugated to the relevant antibody (anti-MqsRA; anti-His for His_6_-YefM). Samples were resolved by 4-20% SDS-PAGE, and then the gel was dried, exposed to Phosphorimaging screens for 1-10 days as needed, and scanned with a Typhoon scanner (GE Healthcare). For the experiment comparing MqsA degradation following expression alone or co-expression with its cognate toxin, the toxin ribosome-binding site was changed to that of the antitoxin to increase toxin expression levels. All pulse-chase data presented are the average of at least two independent biological replicates.

### Western blots

Cells were harvested by centrifugation and flash frozen in liquid nitrogen. Pellets were resuspended in 1x Laemmli buffer and analyzed by SDS-PAGE. Anti-MqsRA antibody was generated using His_6_-MqsRA complex (Covance) and used unpurified at 1:3000. Anti-His_6_ antibody was used at 1:5000 (Thermo Fisher); and purified anti-Lon antibody was used at 1:5000 (kind gift from T. Baker and R. Sauer). SuperSignal West Femto Maximum Sensitivity Substrate (ThermoFisher) was used to develop the blots, which were visualized with a FluorChem R Imager (ProteinSimple). Because high amounts of lysate were loaded onto gels to allow for antitoxin detection, typical loading controls were outside the linear range. Loading normalization was instead done by Coomassie staining membranes after Western blotting. All Western blot data presented are the average of three independent biological replicates.

### Arabinose promoter shut-off experiments

Cells were grown to OD_600_ ∼ 0.3 in M9 media prepared as described previously, but supplemented with 0.4% glycerol instead of glucose. Cells were then induced with 0.4% L-arabinose for 15 min after which they were washed in M9-glucose, and diluted to OD_600_ of 0.01 in M9-glucose. Growth curves were generated on a plate reader as described above.

---

**Table S1.**
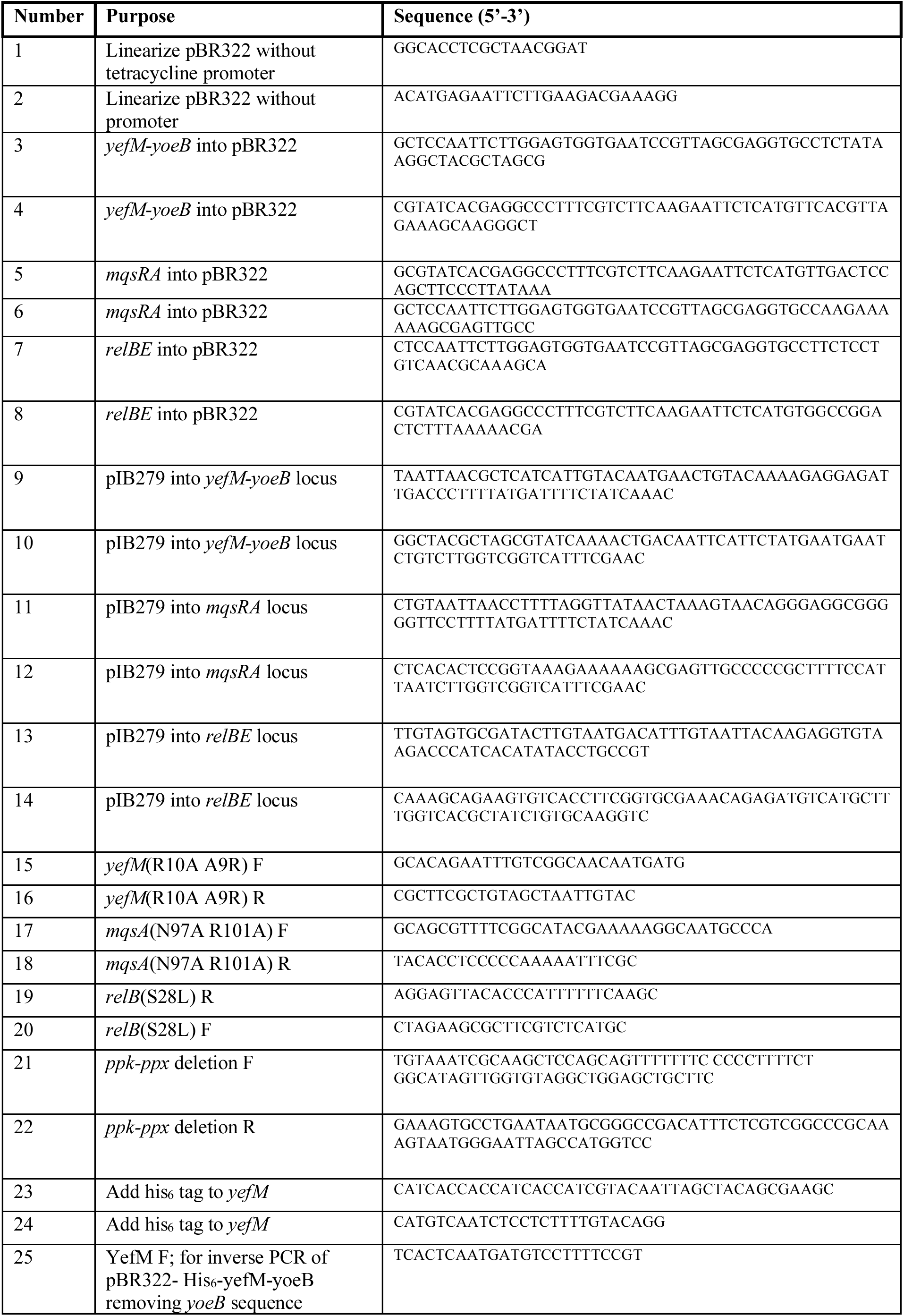
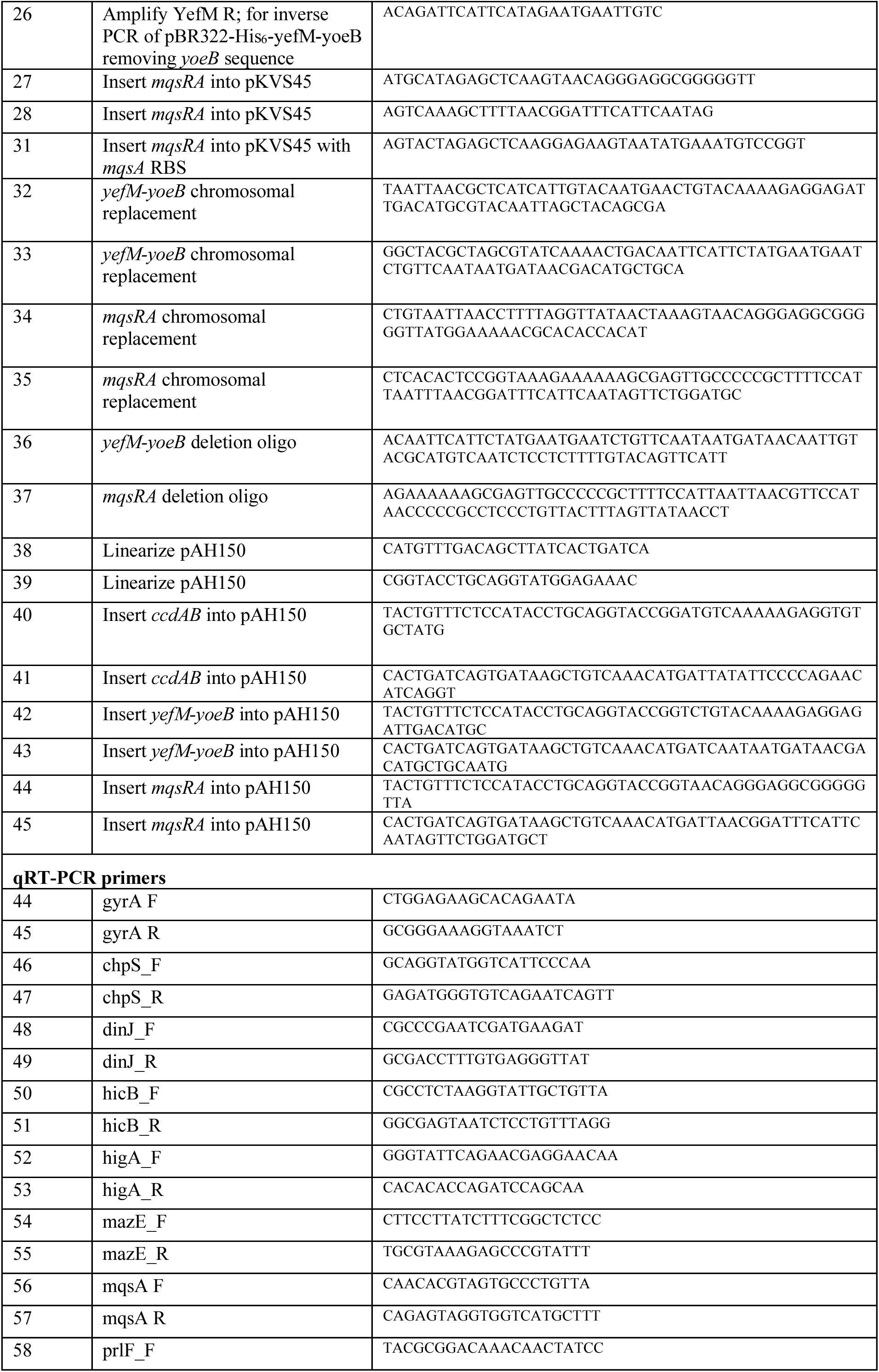
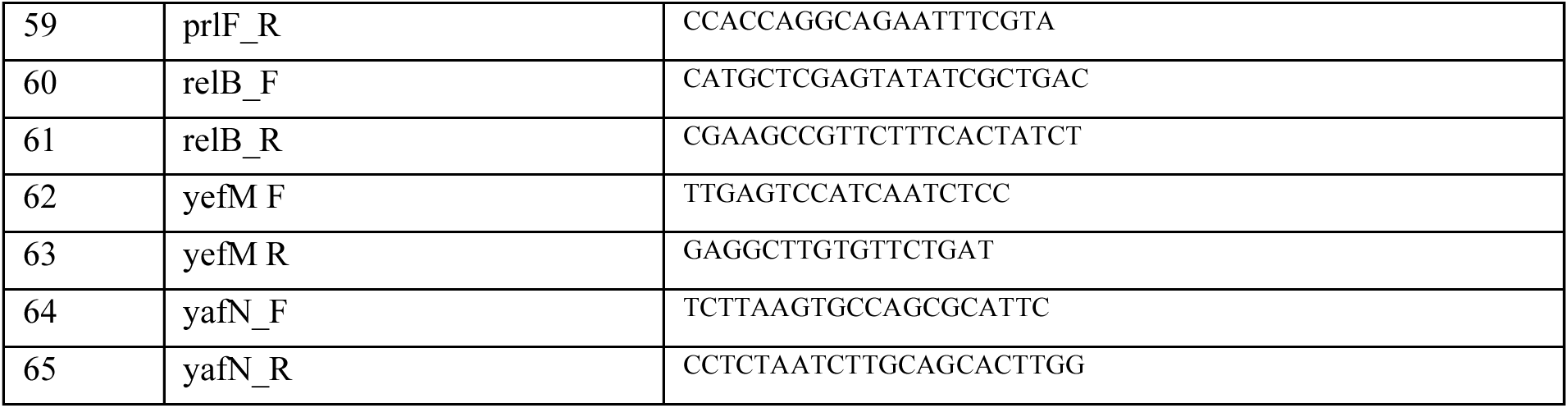
Primers used in this study

**Table S2.** Strains used in this study.

| Name | Genotype | Reference |
| --- | --- | --- |
| ML6 | MG1655 |  |
| ML3203 | *dnaK::kan* | This study |
| ML3204 | *mqsA*<sup>N97A R101A</sup> | This study |
| ML3205 | *yefM*<sup>R10A</sup> | This study |
| ML3206 | *relB*<sup>S28L</sup> | This study |
| ML3207 | $\Delta lon$ | This study |
| ML3208 | $\Delta clpP$ | This study |
| ML3209 | *degP::kan* | This study |
| ML3210 | *hslV::kan* | This study |
| ML3211 | *htpX::kan* | This study |
| ML3212 | *sfhC21* zad220::Tn10 | (Ogura et al., 1999) |
| ML3213 | *sfhC21* zad220::Tn10 $\Delta ftshH3::kan$ | (Ogura et al., 1999) |
| ML3214 | $\Delta ppk \Delta ppx$ | This study |
| ML3215 | pBR322-his<sub>6</sub>-yefM-yoeB | This study |
| ML3216 | $\Delta yefM$-yoeB pBR322-his<sub>6</sub>-yefM(R10A)-yoeB | This study |
| ML3217 | pBR322-his<sub>6</sub>-yefM | This study |
| ML3218 | pBR322-mqsRA | This study |
| ML3219 | pKVS45-mqsRA | This study |
| ML3220 | pKVS45-mqsA | This study |
| ML3221 | MG1655 | (Goeders et al., 2013) |
| ML3222 | $\Delta mazEF \Delta relBE \Delta chpB \Delta dinJ$-yafQ $\Delta yefM$-yoeB<br>$\Delta yafNO \Delta prlF$-yhaV $\Delta hicAB \Delta ygjMN \Delta mqsRA$ | (Goeders et al., 2013) |
| ML3223 | *attL::P<sub>ara</sub>-ccdAB* | This study |
| ML3224 | $\Delta yefM$-yoeB *attL::P<sub>ara</sub>-yefM-yoeB* | This study |
| ML3225 | $\Delta mqsRA$ *attL::P<sub>ara</sub>-mqsRA* | This study |
| ML3264 | *P<sub>lac</sub>-cfp* | This study |
| ML3265 | $\Delta 10TA$ *P<sub>lac</sub>-cfp* | This study |
| ML3266 | *P<sub>lac</sub>-yfp* | This study |
| ML3267 | $\Delta 10TA$ *P<sub>lac</sub>-yfp* | This study |

